# In vivo transomic analyses of glucose-responsive metabolism in skeletal muscle reveal core differences between the healthy and obese states

**DOI:** 10.1101/2022.03.27.486003

**Authors:** Toshiya Kokaji, Miki Eto, Atsushi Hatano, Katsuyuki Yugi, Keigo Morita, Satoshi Ohno, Masashi Fujii, Ken-ichi Hironaka, Yuki Ito, Riku Egami, Saori Uematsu, Akira Terakawa, Yifei Pan, Hideki Maehara, Dongzi Li, Yunfan Bai, Takaho Tsuchiya, Haruka Ozaki, Hiroshi Inoue, Hiroyuki Kubota, Yutaka Suzuki, Akiyoshi Hirayama, Tomoyoshi Soga, Shinya Kuroda

**Affiliations:** Department of Biological Sciences, Graduate School of Science, University of Tokyo, 7-3-1 Hongo, Bunkyo-ku, Tokyo 113-0033, Japan; Data Science Center, Nara Institute of Science and Technology, 8916-5 Takayama, Ikoma, Nara, Japan; Laboratory for Integrated Cellular Systems, RIKEN Center for Integrative Medical Science, 1-7-22 Suehiro-cho, Tsurumi-ku, Yokohama, Kanagawa 230-0045, Japan; Department of Omics and Systems Biology, Niigata University Graduate School of Medical and Dental Sciences, 757 Ichibancho, Asahimachi-dori, Chuo Ward, Niigata City 951-8510, Japan; Institute for Advanced Biosciences, Keio University, Fujisawa, 252-8520, Japan; PRESTO, Japan Science and Technology Agency, 1-7-22 Suehiro-cho, Tsurumi-ku, Yokohama, Kanagawa 230-0045, Japan; Molecular Genetics Research Laboratory, Graduate School of Science, University of Tokyo, 7-3-1 Hongo, Bunkyo-ku, Tokyo 113-0033, Japan; Department of Mathematical and Life Sciences, Graduate School of Integrated Sciences for Life, Hiroshima University 1-3-1 Kagamiyama, Higashi-hiroshima city, Hiroshima, 739-8526, Japan; Department of Computational Biology and Medical Sciences, Graduate School of Frontier Sciences, University of Tokyo, 5-1-5 Kashiwanoha, Kashiwa, Chiba 277-8562, Japan; Division of Integrated Omics, Research Center for Transomics Medicine, Medical Institute of Bioregulation, Kyushu University, 3-1-1 Maidashi, Higashi-ku, Fukuoka 812-8582, Japan; Bioinformatics Laboratory, Faculty of Medicine, University of Tsukuba, Ibaraki, 305-8575, Japan; Center for Artificial Intelligence Research, University of Tsukuba, Ibaraki, 305-8577, Japan; Metabolism and Nutrition Research Unit, Institute for Frontier Science Initiative, Kanazawa University, 13-1 Takaramachi, Kanazawa, Ishikawa, 920-8641, Japan; Institute for Advanced Biosciences, Keio University, 246-2 Mizukami, Kakuganji, Tsuruoka, Yamagata 997-0052, Japan; Core Research for Evolutional Science and Technology (CREST), Japan Science and Technology Agency, Bunkyo-ku, Tokyo 113-0033, Japan

## Abstract

Metabolic regulation in skeletal muscle is essential for blood glucose homeostasis. Obesity causes insulin resistance in skeletal muscle, leading to hyperglycemia and type 2 diabetes. In this study, we performed multiomic analysis of the skeletal muscle of wild-type (WT) and genetically obese (*ob*/*ob*) mice, and constructed regulatory transomic networks for metabolism after oral glucose administration. Our network revealed that metabolic regulation by glucose-responsive metabolites had a major effect on WT mice, especially carbohydrate metabolic pathways. By contrast, in *ob*/*ob* mice, much of the metabolic regulation by glucose-responsive metabolites was lost and metabolic regulation by glucose-responsive genes was largely increased, especially in carbohydrate and lipid metabolic pathways. We present some characteristic metabolic regulatory pathways found in central carbon, branched amino acids, and ketone body metabolism. Our transomic analysis will provide insights into how skeletal muscle responds to changes in blood glucose and how it fails to respond in obesity.

## Introduction

Blood glucose level is regulated by the cooperative function of many tissues. Insulin, the hormone for lowering blood glucose level, is secreted by pancreatic beta cells when blood glucose level rises. Insulin lowers blood glucose level by stimulating glucose disposal in the skeletal muscle and adipose tissue, and inhibits gluconeogenesis in the liver (Evans et al., 2004). Type 2 diabetes mellitus (T2DM) is one of the most devastating results of obesity, and is characterized by insulin resistance and hyperglycemia (Kahn et al., 2006). Reduced responsiveness of skeletal muscle to insulin is one of the critical aspects of T2DM development (DeFronzo and Tripathy, 2009). T2DM is a multifactorial disease involving many complex signaling pathways in different tissues; thus, a comprehensive analysis might help further our understanding of the molecular mechanisms of this disease.

Metabolism is a series of chemical reactions that convert starting materials into molecules that maintain the living state of cells and organisms. Metabolic reactions, defined as chemical reactions of metabolism, are regulated by metabolic enzymes and metabolites. Metabolic enzymes mainly regulate metabolic reactions at the gene expression level, which is determined by transcription factors; and at the enzyme activity level, which is regulated by post-translational modifications such as phosphorylation. Metabolites regulate metabolic reactions through the concentration of substrates, and also through the allosteric regulation of enzyme activity.

Integrating multiple omics techniques such as metabolomics, proteomics, and transcriptomics is useful for understanding the flow of biological information, and has been applied to a wide range of biological problems (Hasin et al., 2017; Wiley, 2011). Several groups have used multiomic approaches to study the molecular mechanisms of insulin resistance. One study integrated epigenomics, transcriptomics, proteomics, and metabolomics to analyze the liver of mice fed a high-fat diet (Soltis et al., 2017). Another study used transcriptomics, proteomics, metabolomics, and microbiomics to analyze blood and stool samples from healthy human participants during weight gain and weight loss (Piening et al., 2018). A transomic approach, proposed by our group, connects measurements of multiple omics layers such as proteomics, transcriptomics, and metabolomics based on direct molecular interactions (Kawata et al., 2018; Yugi and Kuroda, 2018; Yugi et al., 2014, 2016). This approach provides an understanding of the spatiotemporal dynamics of the biochemical network.

We previously performed a transomic study of glucose-responsive molecules in the livers of wild-type (WT) and genetically obese mice (*ob/ob* mice) during oral glucose administration (Kokaji et al., 2020), and an inter-organ transomic study using the liver and skeletal muscle of WT and *ob/ob* mice in the starved state (Egami et al., 2021). In this study, we performed transomic analysis, including transcriptomics and metabolomics, of glucose-responsive molecules in the skeletal muscle of WT and *ob/ob* mice during oral glucose administration. By analyzing time-series data, we identified pathways that are activated or inhibited by oral glucose administration, and determined how they are dysregulated in obesity. Our study provides a better understanding of the mechanism of glucose metabolism in skeletal muscle and T2DM.

## Results

### Overview of the study

Metabolic reactions, which are defined as chemical reactions of metabolism, are regulated by an integrated network of metabolites as allosteric regulators, substrates, and products; metabolic enzymes; transcription factors; and signaling molecules. To elucidate the regulatory network controlling glucose-responsive metabolic reactions in skeletal muscle, we constructed a regulatory transomic network by integrating metabolic reactions with metabolites, gene expression of metabolic enzymes, and transcription factors, using skeletal muscle excised from C57BL/6J WT mice or *ob/ob* mice at different time points after glucose administration (Fig. S1). The transomic network of the skeletal muscle was constructed according to our previous study of the liver (Kokaji et al., 2020).

Glucose was administered orally to 16 h-fasted WT and *ob/ob* mice, and the gastrocnemius muscle and blood were collected at 0, 20, 60, 120, and 240 min after glucose administration (Fig. 1A). The *ob/ob* mice showed elevated levels of blood glucose and insulin compared to WT mice throughout the study (Fig. S2A). The blood and skeletal muscle data in the fasting state were obtained from our previous studies (Egami et al., 2021; Kokaji et al., 2020). The skeletal muscle data after oral glucose administration were newly obtained in this study (Fig. S2B).

**Fig 1.**
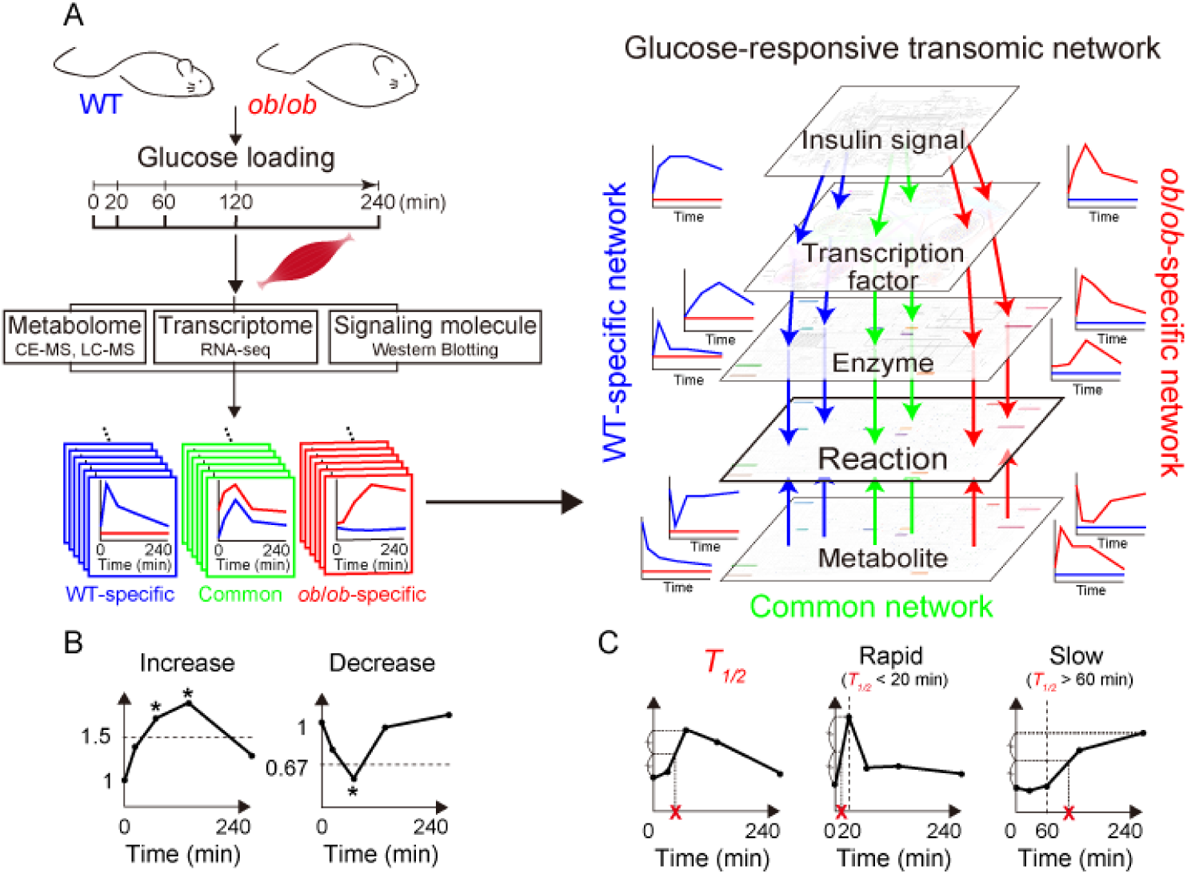
Pipeline of the construction of the glucose-responsive transomic network. **(A)** We measured the time courses of multiomic data from the skeletal muscles of WT and *ob*/*ob* mice following oral glucose administration and identified the molecules that were changed by oral glucose administration, which we defined as glucose-responsive molecules in each layer. We added interlayer regulatory connections between glucose-responsive molecules in different layers using bioinformatics methods and information in public databases. The result was a glucose-responsive transomic network in the skeletal muscle of WT and *ob*/*ob* mice. **(B)** Definition of glucose-responsive molecules using fold change and FDR-adjusted p value. **(C)** Definition of *T_1/2_*, an index of the temporal rate of response, and rapid and slow glucose-responsive molecules using *T_1/2_*.

Using the skeletal muscle data during oral glucose administration, we defined the features of glucose-responsive molecules according to our previous study (Kokaji et al., 2020). Molecules that showed statistically significant changes (absolute log_2_ fold change ≥ 0.585 [2^0.585^ = 1.5] and a false discovery rate [FDR]-adjusted p value [q value] ≤ 0.1) at any time point compared to the fasting state after glucose administration were defined as glucose-responsive (Fig. 1B). We also calculated time constants (*T_1/2_*) to study the temporal patterns of glucose-responsive molecules (Fig. 1C). *T_1/2_* was defined as the amount of time needed for the response to reach half of the minimum (decreasing molecules) or maximum (increasing molecules) amplitude. According to the blood insulin concentration, which peaked at about 20 min and decreased to basal level at about 60 min (Fig. S2A), rapid responses were defined as those with *T_1/2_* values less than 20 min, and slow responses were defined as those with values longer than 60 min.

Glucose-responsive molecules were integrated across the omic layers, and the regulatory transomic network was constructed in WT and *ob*/*ob* mice (Fig. 1A). The transomic networks contained layers of insulin signaling molecules (Insulin signal), transcription factors (TF), gene expression and phosphorylation of metabolic enzymes (Enzyme), metabolic reactions (Reaction), and metabolites (Metabolite), and the layers were connected when regulations could be speculated. By comparing the regulatory transomic networks between WT and *ob/ob* mice, we comprehensively evaluated how obesity affects the responses to glucose in skeletal muscle.

### Metabolomics

We first performed metabolomics analysis using capillary electrophoresis–mass spectrometry (CE–MS), liquid chromatography (LC) –MS, and enzyme assays. A total of 104 water-soluble and ionic metabolites including glucose, amino acids, and nucleic acids were measured by CE–MS. Statistical tests were performed to identify the glucose-responsive metabolites in WT and *ob/ob* mice (Fig. 2A, B; Data File S1). To define an increase or decrease in time courses with changes in both directions at different time points, the direction of change compared to time 0 at the earliest time point that showed a significant change was used. Metabolites that showed statistically significant increases or decreases in WT or *ob/ob* mice are shown in Figure 2A. The responses were categorized into three groups (rapid, intermediate, or slow) according to their *T_1/2_* values (Fig. 2C).

**Fig. 2.**
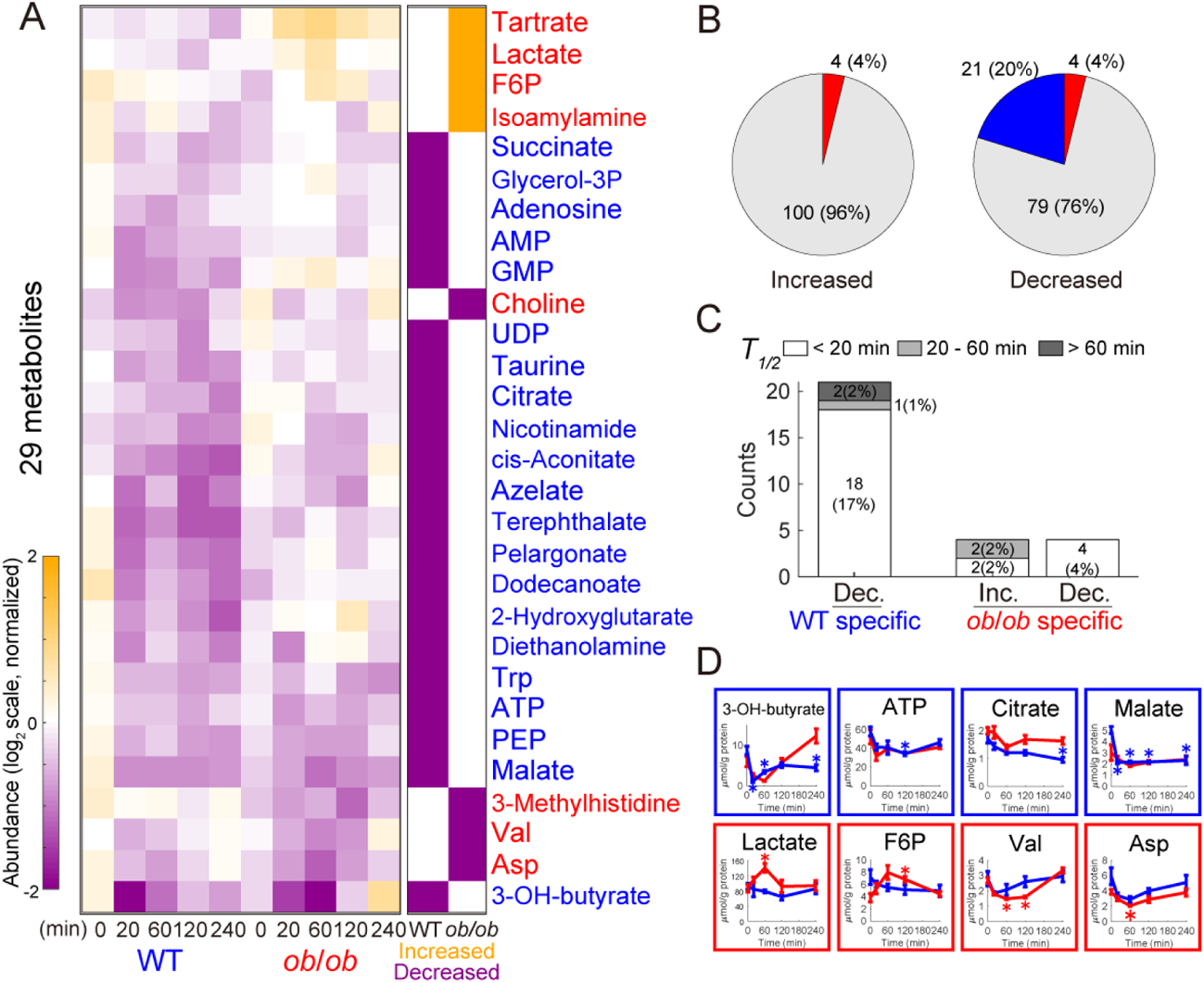
Identification of glucose-responsive metabolites. (**A**) Left: Heat map of the time courses of 29 glucose-responsive metabolites from the skeletal muscles of WT and *ob*/*ob* mice following oral glucose administration. Right: The bars in the heat map are colored according to the extent of glucose responsiveness, meaning the change from fasting state (0 min) in WT and *ob/ob* mice: increased (orange), decreased (purple), or were unchanged (white). Metabolites written in blue text indicate glucose-responsive metabolites specific to WT mice; red text, specific to *ob*/*ob* mice. (**B**) Increased and decreased metabolites in the skeletal muscles of WT mice and *ob*/*ob* mice. Blue, WT specific; red, *ob*/*ob* specific. (**C**) Rapid, intermediate, and slow responses in glucose-responsive metabolites. (**D**) Graphs showing the metabolites with responses that were specific to WT mice (blue boxes) and specific to *ob/ob* mice (red boxes).

Four metabolites (4% of the total quantified metabolites) were significantly increased only in *ob/ob* mice, and none were increased in WT mice (Fig. 2B). Metabolites that were increased only in *ob/ob* mice included fructose 6-phosphate (F6P), tartrate, lactate, and isoamylamine (Fig. 2D). Twenty-one metabolites (20%) were significantly decreased only in WT mice, and four metabolites (4%) were significantly decreased only in *ob/ob* mice (Fig. 2B). It is noteworthy that no common metabolites were increased or decreased in WT and *ob/ob* mice. Metabolites decreased in WT mice included those that play a role in the tricarboxylic acid (TCA) cycle, such as citrate, cis-aconitate, succinate, and malate (Fig. 2D). The ketone body 3-hydroxybutylate (3-OH-butylate) was also decreased in WT mice. Metabolites that were decreased in *ob/ob* mice included valine, aspartic acid, choline, and 3-methylhistidine. Most of the decreased metabolites showed rapid responses in both WT and *ob/ob* mice (Fig. 2C). Hierarchical clustering analysis of the metabolites is shown in Figure S3. LC–MS did not detect significant responses of 14 lipids after oral glucose administration (Data File S2).

Our metabolomic analysis revealed that the number of glucose-responsive metabolites specific to WT mice (21: 0 increased + 21 decreased) was larger than that specific to *ob*/*ob* mice (8: 4 increased + 4 decreased), and no responses were common to both mice. These results indicate that there is a substantial difference in the mechanism of glucose metabolism in skeletal muscle between WT and *ob/ob* mice.

Next, we compared the metabolomic changes in the skeletal muscle and blood. The amount of metabolites was regulated not only within each organ but in the blood circulatory system (Katz and Tayek, 1998). For each metabolite that was measurable in both skeletal muscle and blood (61 metabolites), we calculated the correlation between the time course of the metabolites in the skeletal muscle and that in the blood (Fig. S4A). The blood data were obtained from our previous study (Kokaji et al., 2020). The decreases in 3-OH-butyrate, isoleucine, and leucine were highly correlated between the blood and muscle in WT mice; and the decreases in 3-OH-butyrate and increases in lactate were highly correlated between the blood and muscle in *ob*/*ob* mice (Fig. S4A, B). Our previous study showed that 3-OH-butyrate, isoleucine, and leucine also exhibited a high correlation between the blood and liver in the same mouse (Kokaji et al., 2020). These results suggest that metabolites regulated in the bloodstream are regulated similarly in skeletal muscle and liver.

### Transcriptomics

To elucidate the transcriptional changes and controls in the skeletal muscle of WT and *ob/ob* mice after glucose administration, we performed transcriptomic analysis using RNA sequencing. Of the 14,978 genes analyzed, 4,264 that were significantly changed after oral glucose administration were identified as glucose-responsive genes (Fig. 3A, B; Data File S3). A heatmap of the glucose-responsive genes is shown in Figure 3A. The responses were categorized into three groups (rapid, intermediate, or slow) according to their *T_1/2_*s as in the analysis of the metabolites (Fig. 3C, D). Pathway enrichment analysis was also performed for each type of response (Table 1 and Data File S4). We assigned glucose-responsive genes encoding metabolic enzymes to the Enzyme layer of the transomic network, and glucose-responsive genes encoding transcription factors to the TF layer of the transomic network (Figs. 1 and 5).

**Fig. 3.**
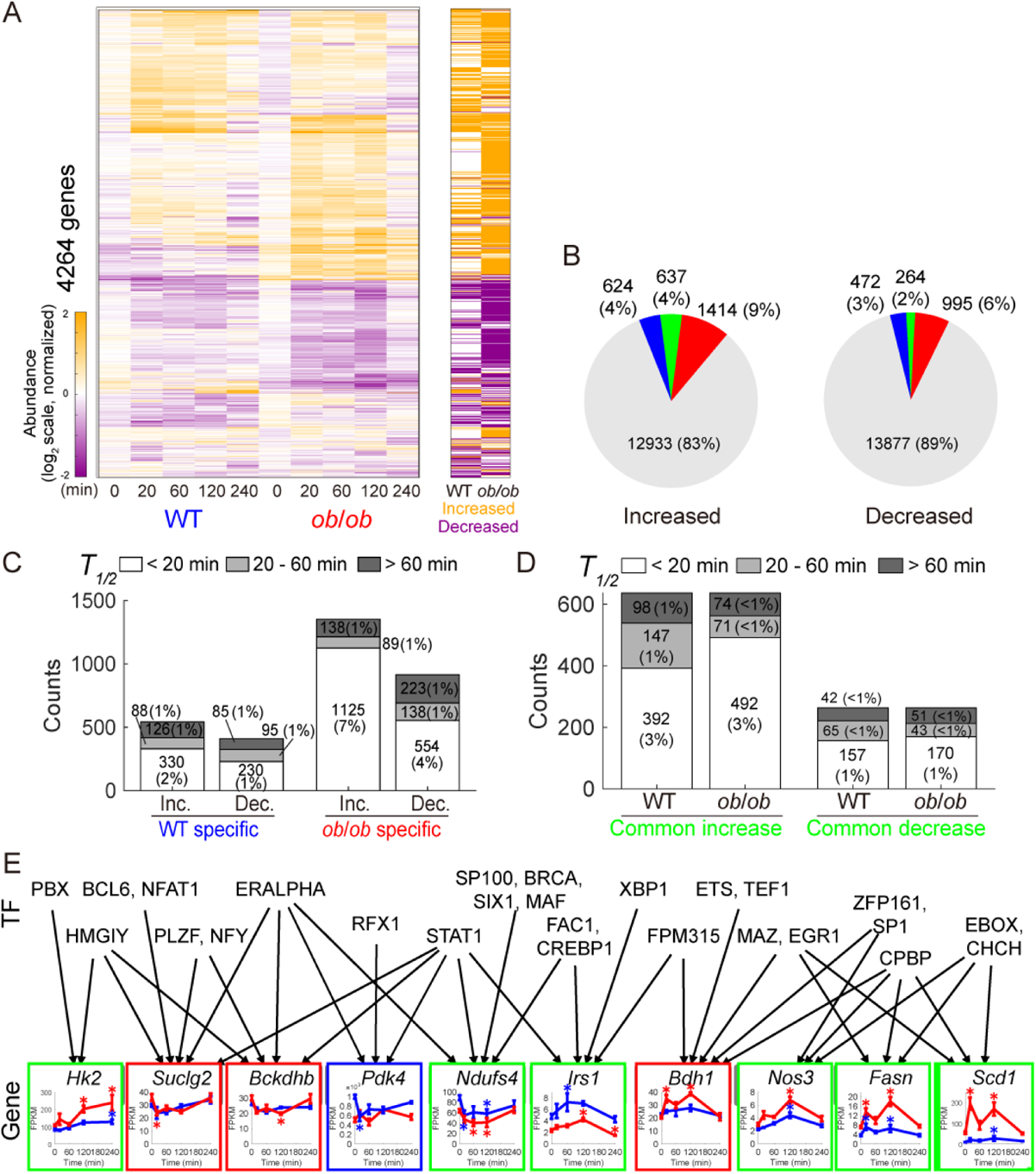
Identification of glucose-responsive genes. (**A**) Left: Heat map of the time courses of transcript abundance for 4,264 glucose-responsive genes in the skeletal muscles of WT and *ob*/*ob* mice. Right: The bars in the heat map are colored according to glucose responsiveness: upregulated (orange) and downregulated (purple). (**B**) Increased and decreased genes in the skeletal muscle of WT mice and *ob*/*ob* mice. Blue, WT specific; red, *ob*/*ob* specific; green, glucose-responsive genes common to both. (**C** and **D**) Rapid, intermediate, and slow responses in glucose-responsive genes. (**E**) Graphs showing the gene expression time courses for the indicated genes. The inferred regulatory connections are shown as arrows from transcription factors to genes.

**Table 1.**
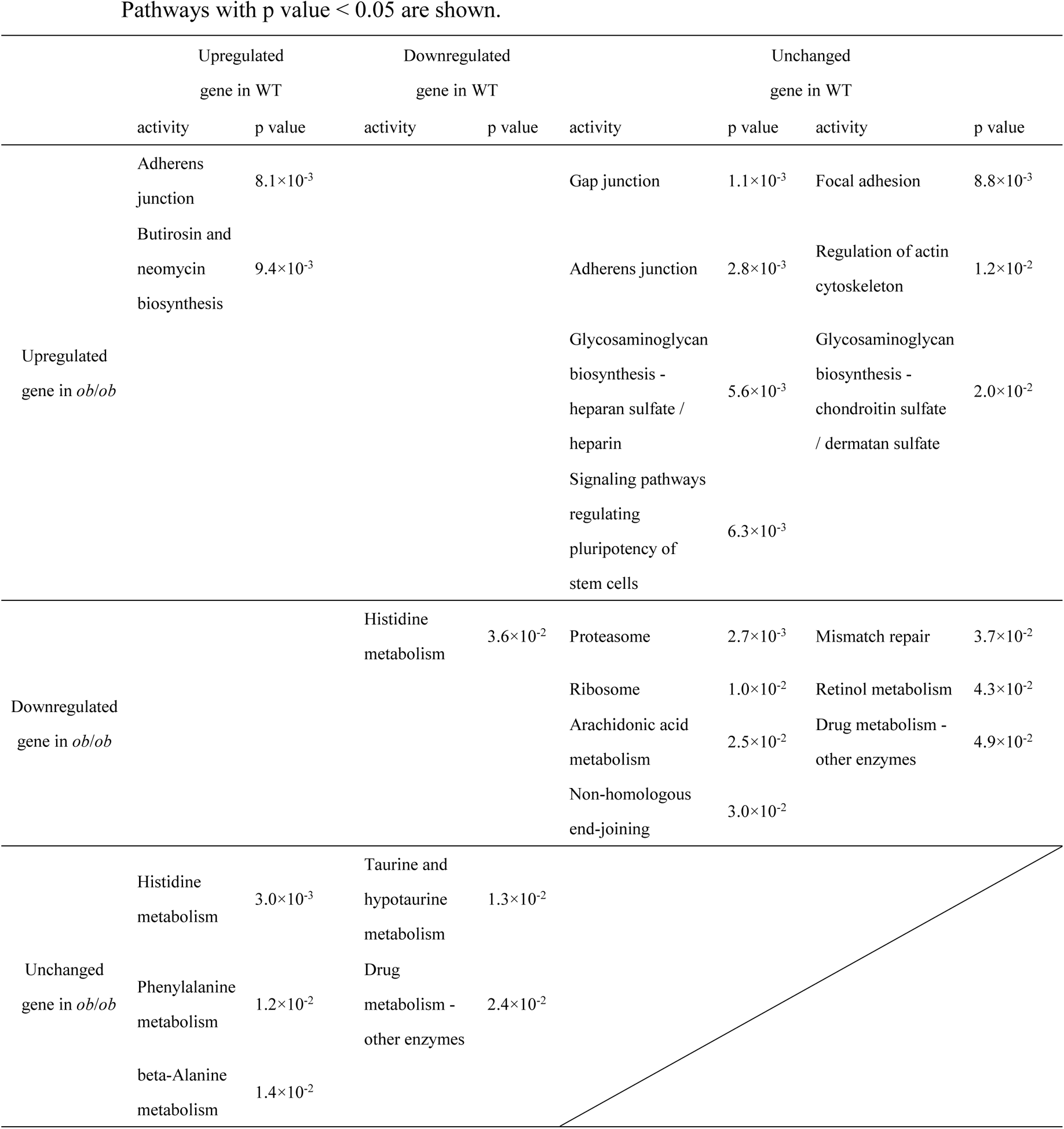
Pathway enrichment analysis of the glucose-responsive genes.

250 Pathways with p value < 0.05 are shown.
|  | Upregulated |  | Downregulated |  | Unchanged |  |  |  |
| --- | --- | --- | --- | --- | --- | --- | --- | --- |
|  | gene in WT |  | gene in WT |  | gene in WT |  |  |  |
|  | activity | p value | activity | p value | activity | p value | activity | p value |
| Upregulated<br>gene in *ob/ob* | Adherens<br>junction | 8.1×10<sup>-3</sup> |  |  | Gap junction | 1.1×10<sup>-3</sup> | Focal adhesion | 8.8×10<sup>-3</sup> |
|  | Butirosin and<br>neomycin<br>biosynthesis | 9.4×10<sup>-3</sup> |  |  | Adherens junction | 2.8×10<sup>-3</sup> | Regulation of actin<br>cytoskeleton | 1.2×10<sup>-2</sup> |
|  |  |  |  |  | Glycosaminoglycan<br>biosynthesis -<br>heparan sulfate /<br>heparin | 5.6×10<sup>-3</sup> | Glycosaminoglycan<br>biosynthesis -<br>chondroitin sulfate<br>/ dermatan sulfate | 2.0×10<sup>-2</sup> |
|  |  |  |  |  | Signaling pathways<br>regulating<br>pluripotency of<br>stem cells | 6.3×10<sup>-3</sup> |  |  |
| Downregulated<br>gene in *ob/ob* |  |  | Histidine<br>metabolism | 3.6×10<sup>-2</sup> | Proteasome | 2.7×10<sup>-3</sup> | Mismatch repair | 3.7×10<sup>-2</sup> |
|  |  |  |  |  | Ribosome | 1.0×10<sup>-2</sup> | Retinol metabolism | 4.3×10<sup>-2</sup> |
|  |  |  |  |  | Arachidonic acid<br>metabolism | 2.5×10<sup>-2</sup> | Drug metabolism -<br>other enzymes | 4.9×10<sup>-2</sup> |
|  |  |  |  |  | Non-homologous<br>end-joining | 3.0×10<sup>-2</sup> |  |  |
| Unchanged<br>gene in *ob/ob* | Histidine<br>metabolism | 3.0×10<sup>-3</sup> | Taurine and<br>hypotaurine<br>metabolism | 1.3×10<sup>-2</sup> |  |  |  |  |
|  | Phenylalanine<br>metabolism | 1.2×10<sup>-2</sup> | Drug<br>metabolism -<br>other enzymes | 2.4×10<sup>-2</sup> |  |  |  |  |
|  | beta-Alanine<br>metabolism | 1.4×10<sup>-2</sup> |  |  |  |  |  |  |

The number of upregulated and downregulated genes in WT and *ob/ob* mice is shown in Figure 3B. The number of glucose-responsive genes specific to *ob/ob* mice (1,414 upregulated, 995 downregulated) was larger than that specific to WT mice (624 upregulated, 472 downregulated). A total of 637 common genes were upregulated and 264 were downregulated in WT and *ob/ob* mice. The calculation of time constants revealed that the number of rapidly responding glucose-responsive genes was larger in *ob/ob* mice than in WT mice (Fig. 3C). Genes upregulated in both WT and *ob/ob* mice included those involved in central carbon metabolism, such as hexokinase 2 (*Hk2*), fatty acid synthase (*Fasn*), and stearoyl-coenzyme A (CoA) desaturase 1(*Scd1*), and the responses in *ob*/*ob* mice were larger than those in WT mice (Fig. 3E). Some genes involved in the insulin signaling pathway also showed upregulation common to both WT and *ob*/*ob* mice, such as insulin receptor substrate 1 (*Irs1*) and nitric oxide synthase 3 (*Nos3*) (Fig. 3E). Genes downregulated in both WT and *ob/ob* mice included those involved in oxidative phosphorylation such as NADH dehydrogenase (ubiquinone) iron-sulfur protein 4 (*Ndufs4*) (Fig. 3E). Genes specifically downregulated in WT mice contained pyruvate dehydrogenase kinase 4 (*Pdk4*) (Fig. 3E). Genes specifically upregulated in *ob/ob* mice were relatively enriched in pathways related to cell adhesion (Table 1). The gene 3-hydroxybutyrate dehydrogenase 1 (*Bdh1*), which is involved in ketone body metabolism, was also specifically upregulated in *ob*/*ob* mice. Genes specifically downregulated in *ob/ob* mice included those involved in the TCA cycle such as succinyl-CoA synthetase beta subunit (*Suclg2*), and those involved in branched-chain amino acid (BCAA) degradation such as 2-oxoisovalerate dehydrogenase beta subunit (*Bckdhb*) (Fig. 3E). Genes specifically downregulated in *ob/ob* mice were relatively enriched in the proteasome pathway and ribosomal proteins (Table 1).

Next, we performed hierarchical clustering analysis of transcriptome data and bioinformatics analysis of the binding motifs of gene clusters using the transcription factor database TRANSFAC (Figs. 3E and S5A, B; Data Files S5 and S6) to estimate the regulatory connections between transcription factors and genes (Kel et al., 2003; Matys et al., 2006). We predicted the regulatory connections between a transcription factor and a gene if the binding motifs of the transcription factor were enriched in the promoter regions of the genes in a cluster. For example, we inferred that early growth response protein 1 (Egr1) is a transcription factor that regulates some of the genes upregulated in WT and *ob/ob* mice (Fig. 3E). A comparison of the estimated regulatory connections with those predicted from chromatin immunoprecipitation (ChIP) experimental data from the ChIP-Atlas database (http://chip-atlas.org/) (Oki et al., 2018) showed that the results from the two methods mostly overlapped (Fig. S5C; Data File S7). The estimated regulatory connections between the transcription factors and the genes encoding metabolic enzymes acted as connections between the TF layer and the Enzyme layer in the transomic network.

### Phosphorylation of insulin signaling molecules

Phosphorylation is an important factor for regulating metabolic reactions. Direct phosphorylation of an enzyme can regulate its activity, and phosphorylation of a transcription factor can regulate the expression level of downstream enzymes. Therefore, we measured the phosphorylation of 10 enzymes, transcription factors, and signaling molecules in the insulin pathway by performing western blot analysis of protein samples prepared from the skeletal muscle of WT and *ob*/*ob* mice during oral glucose administration (Fig. S6; Data File S8). The band intensities were quantified, and the results were used to determine if the phosphorylation was glucose-responsive.

We were able to detect many glucose-responsive phosphorylated proteins from the analysis (Fig. 4). The level of phosphorylated ribosomal protein S6 was increased in both WT and *ob/ob* mice. The phosphorylation of Akt was specifically increased in WT mice, and the phosphorylation of glycogen phosphorylase was specifically decreased in WT mice. Glycogen synthase kinase 3 β (Gsk3β) and cAMP response element-binding protein (Creb) were specifically increased in *ob/ob* mice. Some molecules showed the opposite responses in WT and *ob/ob* mice. For example, the phosphorylation of forkhead box protein 1 (Foxo1) was transiently increased in WT mice but decreased in *ob/ob* mice; the phosphorylation of glycogen synthase (Gs) was decreased in WT mice and increased in *ob/ob* mice. The phosphorylation of extracellular signal-related kinase (Erk) and AMP-activated protein kinase α (Ampkα) was not affected by glucose administration in both WT and *ob/ob* mice. In the subsequent transomic analysis, metabolic enzymes with glucose-responsive phosphorylation were assigned to the Enzyme layer, and transcription factors with glucose-responsive phosphorylation were assigned to the TF layer.

**Fig. 4.**
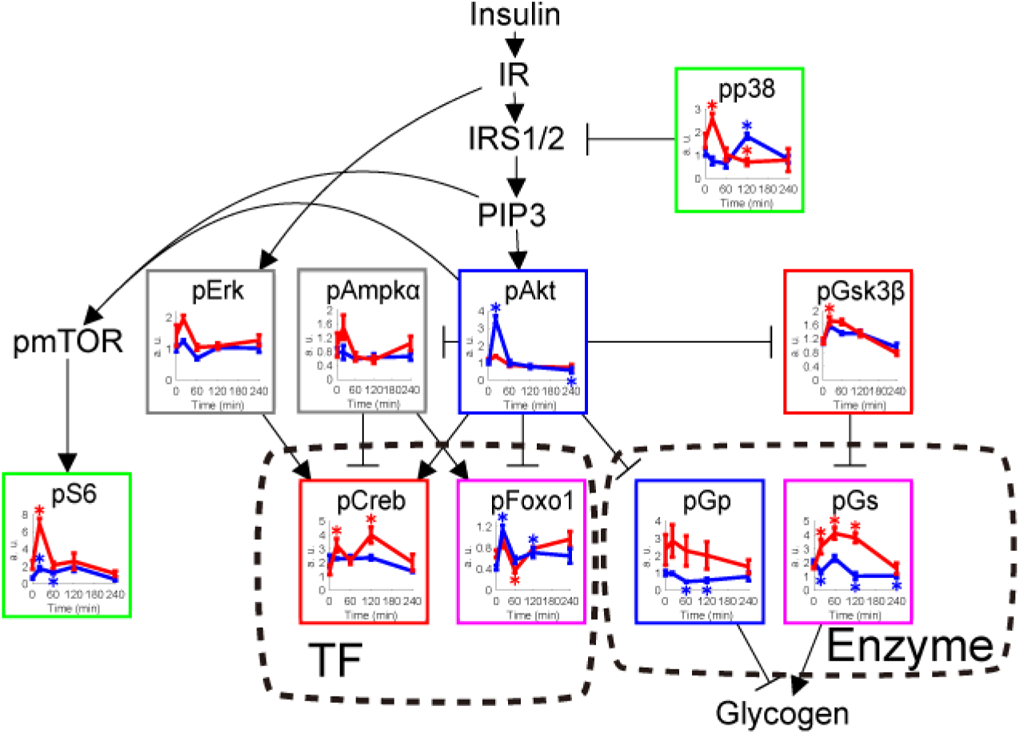
Identification of glucose-responsive phosphorylation of insulin signaling molecules. Time courses of the phosphorylation of the indicated insulin signaling molecules in the skeletal muscle of WT mice (blue lines) and *ob*/*ob* mice (red lines) following oral glucose administration. Phosphorylated proteins are indicated by the prefix “p.” The time course graphs are presented in the context of the insulin signaling pathway from the KEGG database (Kanehisa et al., 2012, 2017). The colors of the boxes around each graph indicate the change in phosphorylation specific to WT (blue), specific to *ob*/*ob* (red), common to both (green), opposite between WT and *ob*/*ob* mice (pink). Proteins that did not exhibit a change in phosphorylation are outlined in gray. Glucose-responsive molecules in the TF and Enzyme layers are enclosed in dashed boxes.

### Regulatory glucose-responsive transomic network

A regulatory transomic network of glucose-responsive molecules in the skeletal muscle was constructed with five layers: Insulin signal, TF, Enzyme, Reaction, and Metabolite (Fig. 5; Data File S9). We constructed the transomic network in the skeletal muscle using a method we previously developed for the transomic network in the liver (Kokaji et al., 2020). Briefly, glucose-responsive molecules were assigned to the corresponding layers as nodes, and the edges between the nodes were drawn to show the interlayer regulatory connections of glucose-responsive molecules retrieved from pathway databases such as Kyoto Encyclopedia of Genes and Genomes (KEGG) and Braunschweig Enzyme Database (BRENDA) (Kanehisa et al., 2012, 2017; Schomburg et al., 2013) (Fig. 5A).

**Fig. 5.**
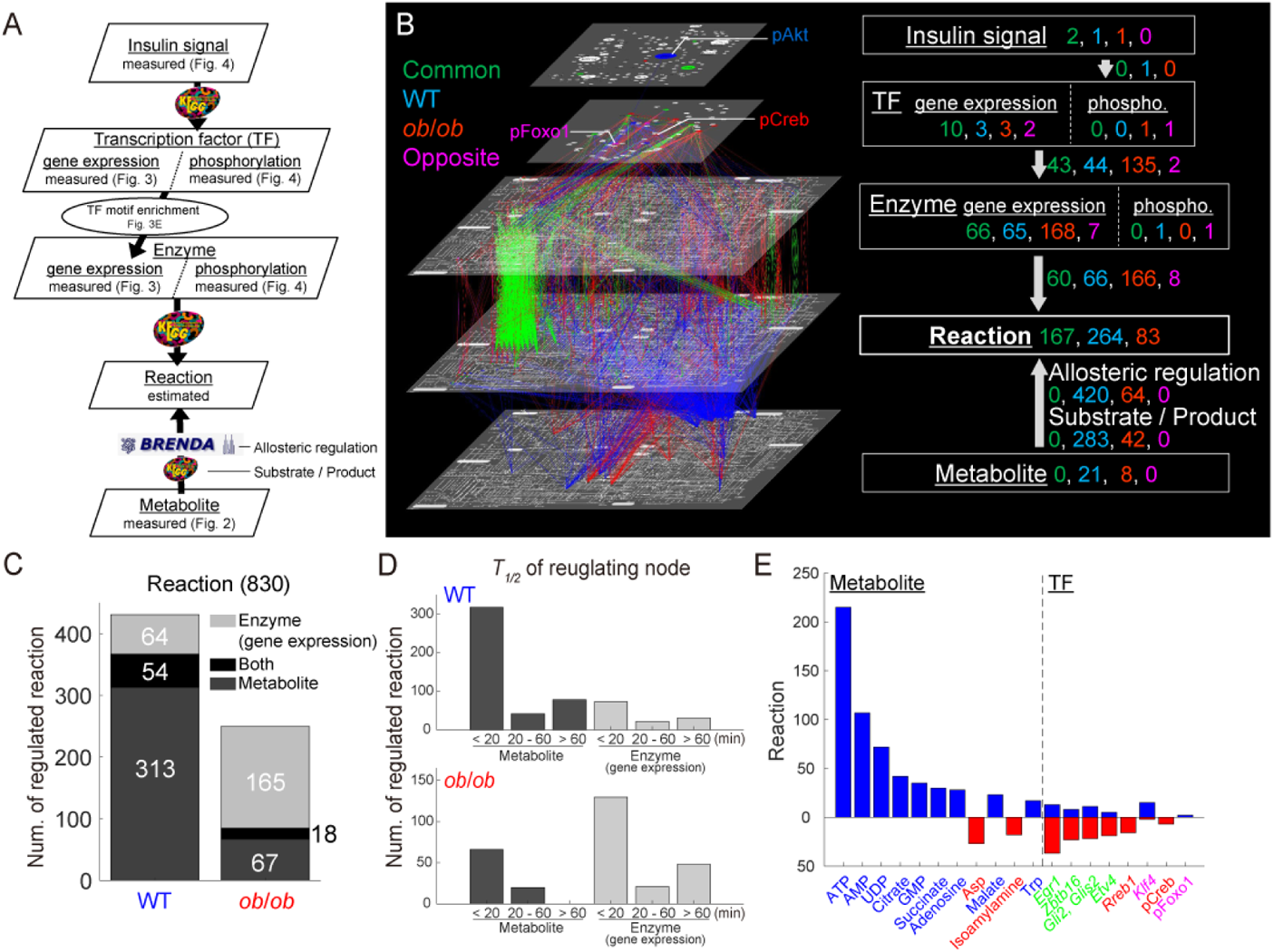
Construction of a regulatory transomic network for glucose-responsive metabolic reactions. (**A**) The procedure for constructing the regulatory transomic network for glucose-responsive metabolic reactions. The databases used to identify the interlayer regulatory connections are shown by arrows. (**B**) The regulatory transomic network for glucose-responsive metabolic reactions. (**C**) The number of glucose-responsive metabolic reactions regulated by glucose-responsive molecules in the Enzyme layer, Metabolite layer, or both. (**D**) The number of glucose-responsive metabolic reactions regulated by glucose-responsive metabolites and genes with the indicated time constants *T_1/2_* in WT mice and *ob*/*ob* mice. (**E**) The number of glucose-responsive metabolic reactions regulated by the indicated glucose-responsive molecules in WT mice (upper, blue) and *ob*/*ob* mice (lower, red). The colors of the names of molecules indicate the type of glucose-responsive molecules as described in (B).

By constructing regulatory transomic networks in WT and *ob/ob* mice, we were able to identify WT specific, *ob/ob* specific, and common responses of molecules and interlayer regulatory connections to glucose administration (Fig. 5B; green, common; blue, WT specific; red, *ob/ob* specific). In the Metabolite layer, the number of WT mice specific glucose-responsive molecules was larger than *ob/ob* mice specific glucose-responsive molecules, and no molecules responded commonly in WT and *ob*/*ob* mice. Therefore, most of the interlayer regulatory connections between the Metabolite layer and the Reaction layer were specific to WT mice, suggesting that metabolic regulation by a metabolite itself after glucose administration is impaired in obesity. By contrast, approximately 55% of glucose-responsive genes in the Enzyme layer and the interlayer regulatory connections between the Enzyme layer and the Reaction layer were classified as *ob/ob* specific, suggesting that transcriptional regulation compensated for the regulation by metabolites that was lost in obese mice. The number of common glucose-responsive genes in the Enzyme layer and its regulatory connections was approximately 40% of the *ob*/*ob* specific ones.

The numbers of glucose-responsive metabolic reactions regulated by metabolites (Metabolite layer), genes (Enzyme layer), or both were calculated (Fig. 5C). The results suggested that the metabolic reactions in WT mice were mainly regulated by metabolites, and those in *ob/ob* mice were mainly regulated through gene expression. We also classified the regulators of metabolic reactions according to their time constants (*T_1/2_*), and revealed that a large number of metabolic reactions was affected by the rapidly responding (<20 min) metabolites and genes in both the WT and *ob*/*ob* networks (Fig. 5D). Glucose-responsive metabolites specific to WT mice included cofactors such as ATP, AMP, and UDP, which could have a large effect on the Reaction layer (Fig. 5E).

### Comparison of the regulatory transomic networks of WT and *ob/ob* mice

To analyze how each metabolic pathway was regulated in the regulatory transomic networks of WT and *ob/ob* mice, we constructed a simplified transomic network using a method that we previously developed (Kokaji et al., 2020) (Fig. 6A, B; Data File S10). Briefly, we converted the Reaction layer into the Pathway layer by placing metabolic reactions in a specific metabolic pathway into a single metabolic pathway node, according to the KEGG metabolic pathway.

**Fig. 6.**
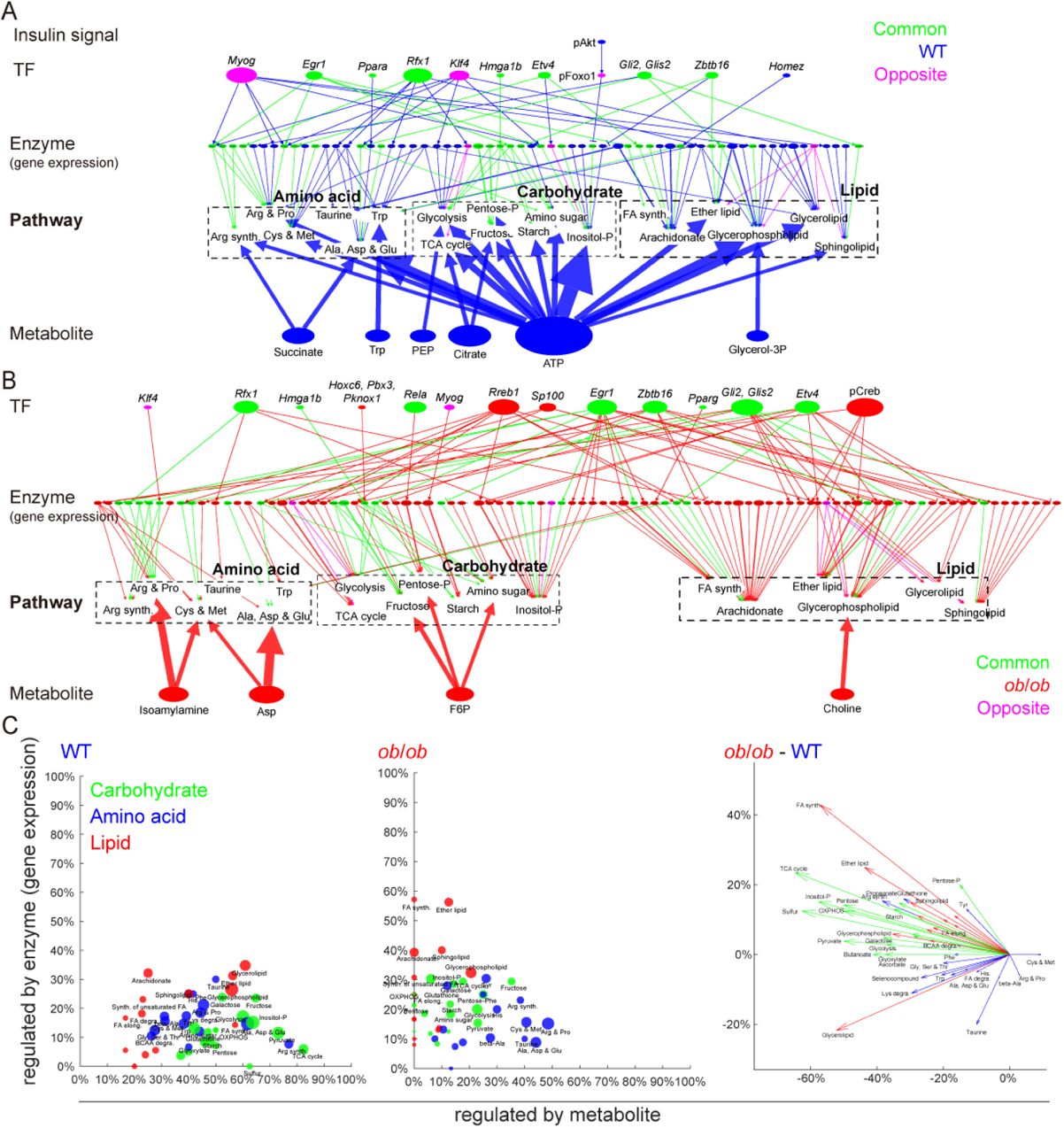
Condensed regulatory transomic networks for glucose-responsive metabolic reactions. (**A, B**) The condensed regulatory transomic network of the response to glucose in WT and *ob/ob* mice. The color of nodes (glucose-responsive molecules) and edges (interlayer regulatory connections) indicate the type of molecules and regulation as described in Figure 5B. The size of the nodes and width of the edges indicate the relative number of the regulated metabolic reactions. (**C**) For each metabolic pathway node, the percentage of regulated metabolic reactions by glucose-responsive metabolites (*x-*axis) and glucose-responsive genes encoding metabolic enzymes (*y-*axis) was plotted for WT and *ob/ob* mice.

In WT mice, various metabolic pathways were regulated by metabolites (Fig. 6A). In particular, carbohydrate metabolic pathways were regulated by WT specific glucose-responsive metabolites such as ATP, citrate, and phosphoenolpyruvate (PEP) (Fig. S7). Although the effects of glucose-responsive genes encoding metabolic enzymes were smaller than the metabolites, some lipid metabolic pathways such as glycerolipid and glycerophospholipid metabolisms were more strongly regulated by glucose-responsive genes than others (Fig. 6C). In *ob*/*ob* mice, the regulation of glucose-responsive metabolites was decreased and that of glucose-responsive genes encoding metabolic enzymes was increased (Fig. 6B). The decreased regulation by metabolites was particularly large in carbohydrate metabolic pathways (Fig. 6C). Regulation by glucose-responsive genes was increased in most carbohydrate and lipid metabolic pathways, with the exception of glycerolipid metabolism. Amino acid metabolic pathways showed relatively small changes in the percentage of metabolic reactions regulated by glucose-responsive metabolites and genes.

**Fig. 7.**
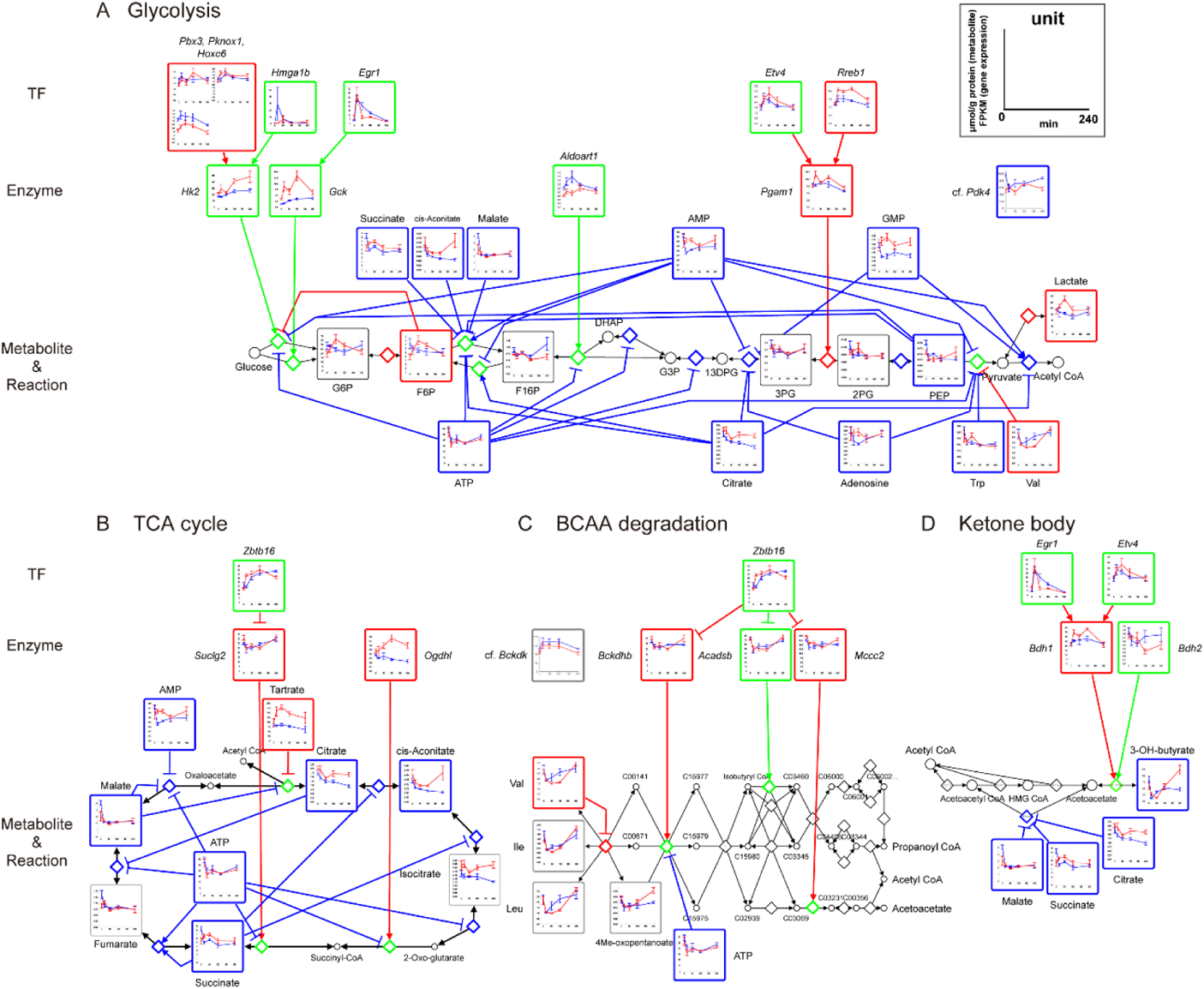
Regulatory transomic network for glucose-responsive metabolic reactions in glycolysis, TCA cycle, BCAA degradation, and ketone body metabolism. The regulatory transomic network for glucose-responsive metabolic reactions in glycolysis (**A**), TCA cycle (**B**), BCAA degradation (**C**), and ketone body metabolism (**D**) in the skeletal muscle of WT mice and *ob*/*ob* mice. Graphs of the time courses of measured molecules are shown for corresponding nodes as the means and SEMs. The colors of the frames and edges indicate WT mice-specific glucose-responsive molecules and regulatory connections (blue), *ob*/*ob* mice-specific glucose-responsive molecules and regulatory connections (red), and common glucose-responsive molecules and regulatory connections and regulatory connections (green). Diamond nodes indicate metabolic reactions.

### Glycolysis, TCA cycle, BCAA degradation, and ketone body metabolism

Finally, we focused on metabolic pathways and their regulatory networks related to glucose (Fig. 7).

#### Glycolysis

In WT mice, although blood glucose levels increased after glucose administration, most metabolites in glycolysis were not defined as “glucose-responsive.” The glycolysis network contained many allosterically regulated WT-specific glucose-responsive metabolites. The decrease in allosteric inhibitors such as ATP and citrate could contribute to the activation of glycolysis in WT mice. We also found upregulation in some glycolytic genes such as *Hk2*, and downregulation in *Pdk4*, which inhibits pyruvate dehydrogenase by phosphorylation (Furuyama et al., 2003). This activation of glycolysis by glucose-responsive molecules might account for the increased influx of glucose from the blood.

In *ob*/*ob* mice, most allosteric regulation was lost, and *Hk2* showed a larger increase than in WT mice, suggesting that the lack of allosteric regulation may be compensated for by gene expression. Because the increase in blood glucose levels was greater than that in WT mice, the increase in F6P and lactate might be caused by an imbalance between increased glucose uptake and activation of glycolytic flux. Glucose 6-phosphate (G6P) was not defined as a glucose-responsive molecule (q value at 60 min = 0.14), but its time series was highly correlated with F6P (Pearson’s r = 0.99). The results are shown in Figure 7A.

#### TCA cycle

In WT mice, four metabolites in the TCA cycle decreased after oral glucose administration (citrate, cis-aconitate, succinate, malate). Although fumarate was not defined as a glucose-responsive molecule (q value at 60 min = 0.13), its time series was highly correlated with malate (Pearson’s r = 0.96). The responses might have caused a decrease in TCA cycle flux and ATP production. The decrease in metabolites in the TCA cycle may be the result of decreased acetyl CoA production derived from β oxidation and ketone body degradation, as well as decreased amino acid degradation and anaplerosis (Dimitriadis et al., 2011a; Furuyama et al., 2003; Puchalska and Crawford, 2017; Saxton and Sabatini, 2017). In *ob*/*ob* mice, the abundance of some metabolites was smaller than that in WT mice before glucose administration, and the metabolites did not show a large response to glucose. The results are shown in Figure 7B. Some studies have reported a decrease in intermediates of the TCA cycle in the skeletal muscle of obese mice (Koves et al., 2008; Wong et al., 2015).

#### BCAA degradation

BCAA degradation pathway and its regulatory network included some glucose-responsive molecules in *ob*/*ob* mice. Valine showed a rapid decrease after oral glucose administration. Leucine and isoleucine were not defined as glucose-responsive molecules (q value at 20 min = 0.15, 0.16), but their time series were highly correlated with valine (Pearson’s r = 0.98 for leucine, 0.98 for isoleucine). The responses may be due to the inhibition of protein degradation by insulin stimulation (Dimitriadis et al., 2011b; Saxton and Sabatini, 2017). Some genes involved in BCAA degradation, such as *Bckdhb*, showed a rapid downregulation. Bckdh kinase (Bckdk) inhibits Bckdh by phosphorylation (Lynch and Adams, 2014), which was not defined as a glucose-responsive molecule (q value at 60 min = 0.11), but its time series was negatively correlated with *Bckdhb* expression (Pearson’s r = −0.96). The transcriptional responses, as well as the decrease in BCAA abundance, might suppress BCAA degradation. We found a similar decrease in *Suclg2* in the TCA cycle, which metabolizes succinyl CoA, one of the BCAA degradation products (Fig. 7B). In WT mice, BCAAs were not defined as glucose-responsive molecules (q value at 20 min = 0.15 to 0.23), but their time series showed a positive correlation with those in *ob*/*ob* mice (Pearson’s r = 0.77 to 0.90) (Fig. 7c).

#### Ketone body metabolism

In WT mice, 3-OH butyrate, a ketone body, showed a rapid and strong decrease (0.13-fold at 20 min). The decrease in metabolites in the TCA cycle, which allosterically inhibit the metabolic enzyme that degrades acetoacetate, might contribute to the degradation of ketone bodies in the skeletal muscle. In *ob*/*ob* mice, 3-OH butyrate did not show a significant decrease (q value at 60 min = 0.12), but *Bdh1* was rapidly upregulated (Fig. 7D).

## Discussion

In this study, we performed transomic analysis of the skeletal muscles obtained from WT and *ob*/*ob* mice after the oral glucose tolerance test to construct a large-scale glucose-responsive regulatory network of metabolism. In WT mice, the number of glucose-responsive metabolites was about 2.5-fold larger than that in *ob*/*ob* mice, and many metabolic reactions were affected by these glucose-responsive metabolites. In particular, the responses of cofactors such as ATP, and TCA cycle intermediates such as citrate and succinate, might affect carbohydrate and amino acid metabolism. By contrast, the number of glucose-responsive genes encoding metabolic enzymes in *ob*/*ob* mice was about 1.8-fold larger than that in WT mice, and the genes were mainly related to carbohydrate and lipid metabolism.

We also found some characteristic glucose-responsive regulatory pathways in central carbon, branched amino acids, and ketone body metabolism. The WT mice showed few significant changes in the metabolites of glycolysis despite the administration of glucose. A recent study showed that the influx of orally administered glucose into the glycolysis of gastrocnemius muscle (white muscle), which was used in this study, is much smaller than that of soleus muscle (red muscle) (Lopes et al., 2021). The decrease in ATP and TCA cycle intermediates also suggested a decrease in TCA cycle flux. Because blood lactate increased (Fig. S4B), much of the glucose imported into the skeletal muscle might be released into the blood as lactate (Brooks, 2020; Hui et al., 2020). In *ob*/*ob* mice, the increase in *Hk2* and F6P indicated an increase in glycolytic flux. In addition, blood lactate increased and TCA cycle intermediates did not respond, suggesting that the conversion of imported glucose to lactate might also occur in *ob*/*ob* mice.

In this study, some amino acids including BCAA in the blood and skeletal muscle were decreased after glucose administration similar to the effect on the liver (Kokaji et al., 2020), suggesting suppression of protein degradation and promotion of protein synthesis in the insulin target organs (Dimitriadis et al., 2011b; Ruvinsky and Meyuhas, 2006). In addition, we found the transcriptional activation of *Bckdk*, a known regulator of the BCAA degradation pathway, and transcriptional repression of the metabolic enzymes, including *Bckdhb*, in *ob*/*ob* mice. These responses might suppress the degradation of amino acids in the skeletal muscle. The blood level of a ketone body, an alternative energy source in the fasting state, was decreased in both WT and *ob*/*ob* mice after glucose administration. We also found that ketone levels in skeletal muscle showed a similar time series as those in the blood, suggesting that intramuscular ketone utilization was also reduced. Decreased degradation of these metabolites could contribute to a decrease in TCA cycle intermediates, but further research is needed to understand why the reduction was specific to WT mice.

We previously constructed a glucose-responsive transomic network in the liver of WT and *ob*/*ob* mice (Kokaji et al., 2020). The liver network contained more glucose-responsive molecules and regulatory connections than the skeletal muscle network, but the differences between WT and *ob*/*ob* mice were similar between the liver and skeletal muscle. In both organs, many metabolic reactions in the WT networks were regulated by metabolites, whereas in the *ob*/*ob* networks, much of the regulation by metabolites was lost and metabolic regulation by gene expression was activated. There were also similarities in the regulation of the metabolic pathway, such as the regulation of carbohydrate metabolism by metabolites and the regulation of lipid metabolism by gene expression. We are currently performing a detailed comparative analysis between the liver network and skeletal muscle network.

To construct a comprehensive glucose-responsive network, it was necessary to integrate more omics data into our network. Because the Insulin signal layer was determined by western blot analysis, the numbers of glucose-responsive molecules and regulatory connections of the layer were very limited compared to those of the other layers. Integration of phosphoproteomic data and kinase-substrate interactions will facilitate a more extensive evaluation of the effects from the Insulin signal layer to the Reaction layer (Humphrey et al., 2013; Krycer et al., 2017; Ohno et al., 2020). The transcription factors of the glucose-responsive genes were determined based on the binding motifs in the promoter sequences and the temporal patterns. Because not all motifs are bound by transcription factors, direct measurements of transcription factor binding using ChIP sequencing analysis will identify a more accurate and extensive regulatory network of glucose-responsive genes (Chèneby et al., 2018; Oki et al., 2018; Yevshin et al., 2019). Although our transomic network was not comprehensive, we revealed several important features of metabolic regulation in the skeletal muscle after glucose administration. An extension of this *in vivo* transomic analysis will lead to a better understanding of glucose homeostasis at the whole-body level and its dysregulation in obesity.

## Materials and Methods

### Animals and sample preparation

Animal experiments were performed as previously described (Kokaji et al., 2020). C57BL/6 WT mice or *ob/ob* mice at ten weeks of age were purchased from Japan SLC Inc. (Shizuoka, Japan). Animal experiments were approved by the animal ethics committee of The University of Tokyo. Overnight-fasted mice were administered an oral glucose load of 2 g/kg body weight. To measure blood glucose and insulin levels, 15 μL blood was collected from the tail veins at 0, 2, 5, 10, 15, 20, 30, 45, 60, 90, 120, 180, and 240 min after glucose administration (n = 5). We used the blood glucose and insulin levels measured in our previous study (Kokaji et al., 2020) (Fig. S2). For the metabolome and transcriptome studies, mice were sacrificed at 0, 20, 60, 120, and 240 min after glucose administration, and the gastrocnemius muscle was excised. Muscle samples were frozen immediately in liquid nitrogen and homogenized with dry ice. The powdered samples were divided and used for metabolomics, lipidomics, transcriptomics, a glycogen assay, and western blotting.

### Metabolomics

Metabolomic analysis was performed as previously described (Kokaji et al., 2020). Total metabolites and proteins were extracted from the skeletal muscle with methanol:chloroform:water (2.5:2.5:1) extraction. Approximately 40 mg of the skeletal muscle was suspended in 500 μL ice-cold methanol containing internal standards (20 μM L-methionine sulfone [Wako, Osaka, Japan], 2-morpholinoethanesulfonic acid, monohydrate [Dojindo, Kumamoto, Japan], and D-camphor-10-sulfonic acid [Wako]) for normalization of MS peak intensities across runs, followed by suspension in 500 μL chloroform, and finally in 200 μL water. After centrifugation at 4,600 × *g* for 15 min at 4°C, the aqueous layer was filtered through a 5 kDa molecular weight cutoff filter (Millipore, Burlington, MA, USA) to remove protein contamination. The filtrate (320 μL) was lyophilized and, prior to MS analysis, dissolved in 50 μL water containing reference compounds (200 μM each of trimesate [Wako] and 3-aminopyrrolidine [Sigma-Aldrich, St. Louis, MO, USA]). Proteins were precipitated by adding 800 μL ice-cold methanol to the interphase and organic layers and centrifuged at 12,000 × *g* for 15 min at 4°C. The pellet was washed with 1 mL ice-cold 80% (v/v) methanol and resuspended in 1 mL sample buffer containing 1% sodium dodecyl sulfate (SDS) and 50 mM Tris-Cl pH8.8, followed by sonication. The total protein concentration was determined by the bicinchoninic acid (BCA) assay and was used for the normalization of metabolite concentration among samples.

All CE–MS experiments were performed using the Agilent 1600 Capillary Electrophoresis system (Agilent Technologies Santa Clara, CA, USA), the G1603A Agilent CE-MS adapter kit, and the G1607A Agilent CE electrospray ionization (ESI) – MS sprayer kit. Briefly, to analyze the cationic compounds, a fused silica capillary (50 µm internal diameter [i.d.] × 100 cm) was used with 1 M formic acid as the electrolyte (Soga and Heiger, 2000). Methanol/water (50% v/v) containing 0.01 µM hexakis(2,2-difluoroethoxy)phosphazene was delivered as the sheath liquid at 10 µL/min. ESI-time-of-flight (TOF) MS was performed in the positive ion mode, and the capillary voltage was set to 4 kV. Automatic recalibration of each acquired spectrum was achieved using the masses of the reference standards ([^13^C isotopic ion of a protonated methanol dimer (2 MeOH+H)]^+^, *m*/*z* 66.0631 and [hexakis(2,2-difluoroethoxy)phosphazene +H]^+^, *m*/*z* 622.0290). To identify the metabolites, the relative migration times of all peaks were calculated by normalization to the reference compound 3-aminopyrrolidine. The metabolites were identified by comparing their *m/z* values and relative migration times to the metabolite standards. Quantification was performed by comparing peak areas to calibration curves generated using internal standardization techniques with methionine sulfone. The other conditions were identical to those previously described (Soga et al., 2006). To analyze anionic metabolites, a commercially available COSMO(+) (chemically coated with cationic polymer) capillary (50 µm i.d. x 105 cm) (Nacalai Tesque, Kyoto, Japan) was used with a 50 mM ammonium acetate solution (pH 8.5) as the electrolyte. Methanol/5 mM ammonium acetate (50% v/v) containing 0.01 µM hexakis(2,2-difluoroethoxy)phosphazene was delivered as the sheath liquid at 10 µL/min. ESI-TOF MS was performed in the negative ion mode, and the capillary voltage was set to 3.5 kV. For anion analysis, trimesate and D-camphor-10-sulfonic acid were used as the reference and internal standard, respectively. The other conditions were identical to those described previously (Soga et al., 2009). Agilent MassHunter software (Agilent technologies) was used for data analysis (Ishii et al., 2007; Soga et al., 2006, 2009).

We used the blood metabolome data obtained in our previous study (Kokaji et al., 2020).

### Lipidomics

Lipidomic analysis was performed as previously described (Egami et al., 2021). Lipidomic profiling of the skeletal muscle was performed by Metabolon, Inc. (Morrisville, NC, USA). Lipids were extracted from samples with dichloromethane and methanol using the modified Bligh and Dyer procedure in the presence of internal standards, with the lower organic phase used for analysis. The extracts were concentrated under nitrogen and reconstituted in 0.25 mL dichloromethane:methanol (50:50) containing 10 mM ammonium acetate. The extracts were placed in vials for infusion–MS analyses, which were performed on the SelexION equipped Sciex 5500 QTRAP mass spectrometer using both the positive and negative ion modes. Each sample was subjected to two analyses, with ion mobility spectrometry–MS conditions optimized for lipid classes monitored in each analysis. The 5500 QTRAP was operated in the multiple reaction monitoring mode to monitor the transitions for more than 1,100 lipids from up to 14 lipid classes. Individual lipid species were quantified based on the ratio of the signal intensity for target compounds to the signal intensity for an assigned internal standard of known concentration. Fourteen lipid class concentrations were calculated from the sum of all molecular species within a class.

### Glycogen assay

Glycogen content was determined as previously described with some modifications (Noguchi et al., 2013). Approximately 20 mg of the skeletal muscle was digested with 1.2 mL of 30% (w/v) potassium hydroxide solution for 1 h at 95°C and neutralized with 61.2 μL glacial acetic acid. The total protein concentration of the muscle digest was determined by the BCA assay and adjusted to 1 μg protein/μL. Glycogen was extracted from the digested skeletal muscle using Bligh and Dyer method to remove lipids (Von Wilamowitz-Moellendorff et al., 2013). The digested skeletal muscle (50 μL) was mixed with 120 μL ice-cold methanol, 50 μL chloroform, 10 μL of 1% (w/v) linear polyacrylamide, and 70 μL water. After incubation on ice for 30 min, the mixture was centrifuged at 12,000 × *g* to remove the separated aqueous layer. The glycogen was precipitated by the addition of 200 μL methanol and centrifugation at 12,000 × *g* for 30 min at 4°C, washed with ice-cold 80% (v/v) methanol, and dried completely. Glycogen pellets were suspended in 20 μL of 0.1 mg/mL amyloglucosidase (Sigma-Aldrich) in 50 mM sodium acetate buffer and incubated for 2 h at 55°C to digest the glycogen. The concentration of the glucose produced from the glycogen was determined using the Amplex Red Glucose/Glucose Oxidase Assay kit (Thermo Fisher Scientific, Waltham, MA, USA), according to the manufacturer’s instructions.

### Transcriptomics

Transcriptomic analysis was performed as previously described (Kokaji et al., 2020). Total RNA was extracted from the skeletal muscle using the RNeasy Mini Kit (QIAGEN, Hilden, Germany) and QIAshredder (QIAGEN); the quantity was assessed using the Nanodrop (Thermo Fisher Scientific) and the quality was assessed using the 2100 Bioanalyzer (Agilent Technologies). cDNA libraries were prepared using the SureSelect strand-specific RNA library preparation kit (Agilent Technologies). The resulting cDNAs were subjected to 100 base paired-end sequencing on the Illumina HiSeq2500 Platform (Illumina, San Diego, CA, USA) (Matsumoto et al., 2014). Sequences were aligned to the mouse reference genome obtained from the Ensembl database (Cunningham et al., 2015; Flicek et al., 2014) (GRCm38/mm10, Ensembl release 97) using the STAR software package (v.2.5.3a) with the parameters “-- quantMode TranscriptomeSAM --outFilterMultimapScoreRange 1 -- outFilterMultimapNmax 20 --outFilterMismatchNmax 10 --alignIntronMax 500000 -- alignMatesGapMax 100000 --sjdbScore 2 --alignSJDBoverhangMin 1 --genomeLoad NoSharedMemory --outFilterMatchNminOverLread 0.33 -- outFilterScoreMinOverLread 0.33 --sjdbOverhang 100 --outSAMattributes NH HI NM MD AS XS --outSAMunmapped Within --outSAMtype BAM SortedByCoordinate -- outSAMheaderHD @HD VN:1.4 --limitBAMsortRAM 103079215104 -- outSAMstrandField intronMotif” (Dobin et al., 2013). The RSEM tool (v.1.3.0) was used to assemble transcript models (Ensembl release 97) from aligned sequences and to estimate gene expression level with the parameters “--estimate-rspd --forward-prob 0.5 - p 12” (Li and Dewey, 2011). Gene expression level was shown as fragments per kilobase of exon per million mapped fragments (FPKM).

### Western blot analysis

Total proteins were extracted from the skeletal muscle with methanol:chloroform:water (2.5:2.5:1). Ice-cold methanol was added to the skeletal muscle at a concentration of 100 mg/mL of the weight of the skeletal muscle, and the suspension (400 μL) was mixed with chloroform (400 μL) and water (160 μL), followed by centrifugation at 4,600 × *g* for 10 min at 4°C. The aqueous and organic phases were removed and 800 μL ice-cold methanol was added to the interphase to precipitate proteins. The resulting pellet was suspended with 400 μL lysis buffer (10 mM Tris-HCl [pH 6.8] in 1% SDS) and incubated for 15 min at 65°C, followed by sonication. The protein lysate was centrifuged at 12,000 × *g* for 3 min at 4°C to remove debris. The total protein concentration of the resulting supernatant was determined by the BCA assay. The following primary antibodies were purchased from Cell Signaling Technology (Danvers, MA, USA): phosphorylated Erk1/2 (p-Erk1/2, Thr^202^/Tyr^204^; #9101), pCreb (Ser^133^; #9198), pAkt (Ser^473^; #9271), pS6 (Ser^235^/Ser^236^; #2211), pGsk3β (Ser^9^; #9336), pGs (Ser^641^; #3891), pFoxo1 (Ser^256^; #9461), pp38 (Thr^180^/Tyr^182^; #9211), and pAmpkα (Thr^172^; #2531); pGp (Ser^15^) was made in house as previously described (Noguchi et al., 2013). The proteins (10 μg) were resolved by SDS-PAGE, electrotransferred to nitrocellulose membranes, and incubated with the appropriate antibodies. Immunodetection was performed using the Immobilon Western Chemiluminescent HRP Substrate (Millipore) or SuperSignal West Pico PLUS Chemiluminescent Substrate (Thermo Fisher Scientific), and the Western blot signals were detected using a luminoimage analyzer (LAS-4000; Fujifilm) and quantified with ImageJ software.

### Identification of glucose-responsive molecules

Glucose-responsive molecules were determined as previously described (Kokaji et al., 2020). Molecules that were detected in less than half of the replicates in either WT or *ob*/*ob* mice at any time point after oral glucose administration were removed from the analysis. A molecule with a statistically significant change in response to oral glucose administration was defined as a glucose-responsive molecule according to the following criteria. The fold change of the mean amount at each time point over the mean amount at fasting state (0 min) was calculated for each molecule. The significance of change at each time point was tested by the two-tailed Welch’s *t*-test for each metabolite and phosphorylation, and by the edgeR package (version 3.26.8) of the R language (version 3.6.1) with the default parameters for each gene (Robinson et al., 2009). Metabolite, gene, and phosphorylation that showed an absolute log_2_ fold change ≥ 0.585 (2^0.585^ = 1.5) and an FDR-adjusted p value (q value) ≤ 0.1 at any time point were defined as a glucose-responsive metabolite (Fig. 2A, B), gene (Fig. 3A, B), and phosphorylation (Fig. 4). The q values were calculated by Storey’s procedure (Storey, 2002). To define an increase or decrease in time courses with changes in both directions at different times, we used the direction of change compared to time 0 at the earliest time point that showed a significant change.

### Clustering analysis

Time courses for each metabolite of WT mice and *ob*/*ob* mice were normalized by dividing by the geometric mean of the values of WT mice and *ob*/*ob* mice in the fasting state (0 min) followed by log_2_ transformation. We combined the two time courses of WT and *ob*/*ob* mice for each metabolite and performed hierarchical clustering of the combined time courses using Euclidean distance and Ward’s method (Fig. S3). Based on the clustering tree, we defined eight different clusters of metabolites, showing similar or different responses between WT and *ob*/*ob* mice.

Clustering analysis of gene expression was performed as previously described with some modifications (Kokaji et al., 2020). Time courses for the expression of each gene of WT and *ob*/*ob* mice were normalized by subtracting the average expression values of the time courses of both mice and then dividing the resulting values by the standard deviation (Z-score normalization). We combined the two time courses of WT and *ob*/*ob* mice for each gene and performed hierarchical clustering of the combined time courses using Euclidean distance and Ward’s method (Fig. S7A). The genes with significant differences between WT and *ob*/*ob* mice before glucose administration (0 min) (q value < 0.1) or a significant response at any time point in either WT or *ob*/*ob* mice (q value < 0.1) were selected for the clustering analysis (12301 genes). For the selection, the p value was calculated using the edgeR package (version 3.26.8) of the R language (version 3.6.1) with the default parameters (Robinson et al., 2009), and the q value was calculated by Storey’s procedure (Storey, 2002).

### Pathway enrichment analysis

We performed pathway enrichment analysis of glucose-responsive genes (Table 1; Data File S4). The enrichment of the genes in each pathway was determined using the one-tailed Fisher’s exact test. We used the genes detected in more than half of the replicates in WT and *ob*/*ob* mice at all time points as background. We used the pathways in Metabolism, Genetic Information Processing, and Cellular Processes from the KEGG database (Kanehisa et al., 2012, 2017).

### Prediction of the transcription factor binding motif and inference of regulatory connections between transcription factors and genes

Analysis of transcription factors was performed as previously described (Kokaji et al., 2020). The flanking regions around the major transcription start site of genes were extracted from GRCm38/mm10 (Ensembl, release 97) using Ensembl BioMart (Kinsella et al., 2011). The region from −300 bp to +100 bp of the major transcription start site was defined as the flanking region, according to FANTOM5 analysis of the time course (Arner et al., 2015). The transcription factor binding motifs in each flanking region (Fig. S5B) were predicted using TRANSFAC Pro, a transcription factor database, and Match, a transcription factor binding motif prediction tool (Kel et al., 2003; Matys et al., 2006). The threshold for each transcription factor binding motif prediction was set using extended vertebrate_non_redundant_min_FP.prf, a parameter set in TRANSFAC Pro (Kokaji et al., 2020).

For the inference of regulatory connections between transcription factors and genes, we performed transcription factor motif enrichment analysis of the genes in each cluster (Fig. S5B). The enrichment of transcription factor binding motif in the flanking regions of genes in each cluster was determined by the one-tailed Fisher’s exact test, and transcription factor binding motifs with q value ≤ 0.1 were defined as significantly enriched. The q values were calculated by the Benjamini–Hochberg procedure (Yoav Benjamini, 1995). We used the genes analyzed in the hierarchical clustering as background. To reduce the number of statistical tests, the clusters that contained ≥ 100 genes were analyzed. If a transcription factor binding motif was enriched in the promoter regions of the genes in a cluster, we inferred the regulatory connections between the corresponding transcription factor and the genes in the cluster. To avoid overestimation, we excluded a cluster from the inference if the transcription factor binding motif was more enriched in the children clusters that contained ≥ 100 genes. To compare the enrichment of transcription factor binding motifs between clusters, we calculated the odds ratio of the transcription factor binding motifs for each cluster.

For validation of the inferred regulatory connections, we examined the overlap between the inferred genes of each transcription factor and those predicted from experimental ChIP data from the ChIP-Atlas database (Oki et al., 2018) (Fig. S5C). The genes for which ChIP sequencing peaks of a transcription factor were detected in the flanking region around the transcription start sites were obtained using “Target Genes,” a prediction tool in the ChIP-Atlas. We used the flanking regions from −1000 bp to +1000 bp of the transcription start sites in Target Genes. The overlap between the inferred genes and genes from ChIP data was determined by the one-tailed Fisher’s exact test, and those with q value ≤ 0.1 were defined as significant. The q values were calculated by the Benjamini–Hochberg procedure (Yoav Benjamini, 1995).

### Insulin signaling pathway

The insulin signaling pathway in Figure 4 is a subset of the nodes of the insulin signaling pathway in the KEGG database (mmu04910) (Kanehisa et al., 2012, 2017). We added regulatory input to Creb from the PI3K-Akt signaling pathway (mmu04151), MAPK signaling pathway (mmu04010), and AMPK signaling pathway (mmu04152), and regulatory input to FoxO1 from the FoxO signaling pathway (mmu04068) in the KEGG database. The edges from Akt to Ampk and from p38 to insulin receptor substrate were added according to previous studies (Archuleta et al., 2009; Jaiswal et al., 2019).

### Construction of the regulatory glucose-responsive transomic network

The transomic network was constructed as previously described with some modifications (Kokaji et al., 2020). The regulatory glucose-responsive transomic networks consisted of five layers, namely Insulin signal, TF, Enzyme, Reaction, and Metabolite, with interlayer regulatory connections (Fig. 5A, B). The Insulin signal layer is the insulin signaling pathway constructed in our previous phosphoproteomic study (Kawata et al., 2018). We included in the Insulin signal layer signaling molecules that we analyzed by western blotting; we did not include transcription factors such as Foxo1, or metabolic enzymes such as Gs in this layer. The TF layer consisted of all transcription factors with an inferred regulatory connection (Fig. S7B). The Enzyme layer consisted of all metabolic enzymes in the pathways in Metabolism obtained from the KEGG database (Kanehisa et al., 2012, 2017). The Reaction layer consisted of the metabolic reactions (based on EC number) corresponding to the metabolic enzymes in the Enzyme layer. The Metabolite layer consisted of all metabolites analyzed by CE–MS. Only the molecules and reactions corresponding to genes that were expressed in at least one sample were included in the Insulin signal, TF, Enzyme, and Reaction layers. Not all 15,608 genes were included in the network.

Glucose-responsive molecules were assigned to the corresponding layers as nodes. The Insulin signal layer consisted of insulin signaling molecules with glucose-responsive phosphorylation. The TF layer consisted of transcription factors encoded by glucose-responsive genes or those with glucose-responsive phosphorylation. The Enzyme layer consisted of metabolic enzymes encoded by glucose-responsive genes or those with glucose-responsive phosphorylation. The Reaction layer consisted of “glucose-responsive metabolic reactions,” which were defined as metabolic reactions regulated by glucose-responsive molecules. The Metabolite layer consisted of glucose-responsive metabolites. We also determined the direction of glucose responsiveness. To determine a direction for time courses with both increased and decreased time points, we used the direction of change at the earliest time point with a significant difference from time 0 (fasting state). We did not determine a direction (increase or decrease) for metabolic reactions because we did not measure metabolic reaction activity.

To determine regulatory connections from the Enzyme and Metabolite layers to the Reaction layer, both the target of the regulatory connection (a metabolic reaction) and the regulating molecule (enzyme or metabolite) had to be glucose-responsive. Among the Insulin signal, TF, and Enzyme layers, the interlayer regulatory connections were determined using the directions of glucose responsiveness of the regulating molecule and the regulated molecules, and the types of interlayer regulatory connections, which were designated as either positive or negative. We defined positive interlayer regulatory connections as when both the regulating molecule and regulated molecule showed the same direction of change, namely, both increased or both decreased. We defined negative interlayer regulatory connections as when the regulating molecule and regulated molecule showed responses in the opposite direction, namely, one increased and the other decreased.

The interlayer regulatory connections between glucose-responsive molecules were determined according to databases. The interlayer connections from the Insulin signal layer to the TF layer were determined by the regulation of transcription factors by kinases retrieved from the KEGG database (Kanehisa et al., 2012, 2017). The interlayer connections from the TF layer to the Enzyme layer were determined from inferred regulatory connections between transcription factors and genes (Fig. 3E). The interlayer connections from the Enzyme layer to the Reaction layer were determined by connecting metabolic reactions to their corresponding metabolic enzymes according to the KEGG database (Kanehisa et al., 2012, 2017). The interlayer connections from the Metabolite layer to the Reaction layer comprised two types of regulatory connections: those mediated by allosteric regulators, which were retrieved from the BRENDA database (Schomburg et al., 2013), and those mediated by the substrate or product of the reaction, which were retrieved from the KEGG database (Kanehisa et al., 2012, 2017). The types of regulatory connections made by glucose-responsive transcription factors were defined according to the Gene Ontology (GO) annotations obtained from the Mouse Genome Database (Bult et al., 2008) (Data File S6). The transcription factors that were included in the list of DNA-binding transcription repressors (GO:0001227) and not in the list of DNA-binding transcription activators (GO:0001228) were defined as transcription repressors. Foxo1 was added to the list of transcription activators based on previous studies of gluconeogenesis (Barthel et al., 2005; Nakae et al., 2001). The effects of the phosphorylation of transcription factors on the types of regulatory connections were defined according to the KEGG database (Kanehisa et al., 2012, 2017). We used the allosteric regulation reported for mammals (*Bos taurus*, *Felis catus*, *Homo sapiens*, “Macaca,” “Mammalia,” “Monkey,” *Mus booduga*, *Mus musculus*, *Rattus norvegicus*, *Rattus rattus*, *Rattus sp*., *Sus scrofa*, “dolphin,” and “hamster”) according to the BRENDA database (Schomburg et al., 2013). Because the reversibility of metabolic reactions was not determined, metabolic reactions were assumed to be regulated by both the substrate and product.

### Generation of a condensed transomic network based on metabolic pathway information

We condensed the regulatory transomic networks as previously described with some modifications (Kokaji et al., 2020). First, we grouped the related metabolic reactions in a specific metabolic pathway into one “metabolic pathway node” (Pathway layer), and classified the metabolic pathway nodes into three classes—carbohydrate, lipid, and amino acid—according to the KEGG database (Kanehisa et al., 2012, 2017). Second, we selected two types of metabolic pathway nodes: one was a pathway that exhibited significant associations with any glucose-responsive metabolites or transcription factors; the other was a pathway whose percentage of regulated reactions was in the top 10% either by glucose-responsive metabolites or by glucose-responsive genes encoding metabolic enzymes (Fig. 6C). The association between the metabolic reactions in a metabolic pathway and those regulated by a glucose-responsive molecule was determined by the one-tailed Fisher’s exact test, and associations with a q value ≤ 0.01 were defined as significant. The q values were calculated by the Benjamini–Hochberg procedure (Yoav Benjamini, 1995). We also selected glucose-responsive metabolites that exhibited significant associations with any metabolic pathway nodes and glucose-responsive transcription factors that regulate five or more metabolic enzymes. Third, we reduced the interlayer regulatory connections from the Metabolite layer to the Pathway layer by removing the interlayer regulatory connections that regulated fewer than five metabolic reactions.

### Implementation

Statistical tests, clustering analysis, enrichment analysis, and transomic network analysis were done using MATLAB 2020a (The Mathworks Inc.). Visualization of transomic network in the Graph Modeling Language formats was done using Python 2.7 and VANTED (Junker et al., 2006).

## Supporting information

Data File S10. Significant associations between glucose-responsive molecules and metabolic pathways.

Data File S1. Metabolomic data.

Data File S2. Lipidomic data.

Data File S3. Transcriptomic data.

Data File S4. Pathway enrichment analysis of glucose-responsive genes.

Data File S5. Enrichment analysis of gene clusters.

Data File S6. Inferred regulatory connections between transcription factors and genes.

Data File S7. Overlap between the inferred genes of transcription factors and those predicted from experimental ChIP data.

Data File S8. Western blotting data.

Data File S9. Regulatory transomic network for glucose-responsive metabolic reactions.

## Supplementary Materials

**Fig. S1.**
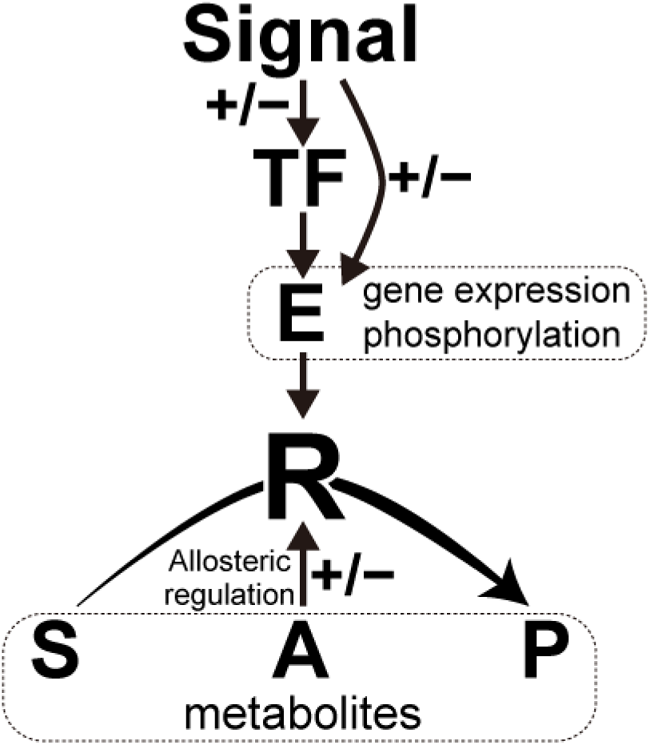
The regulatory network for metabolic reactions. A generic metabolic reaction (R) is catalyzed by a metabolic enzyme (E) and involves metabolites that function as the substrate (S), product (P), or allosteric regulator (A). For reversible reactions, the product is also a substrate and the substrate is also a product (not shown). Positive and negative signs indicate positive and negative regulation, respectively. Regulation of a metabolic reaction by a metabolic enzyme consists of regulation by changing the amount of enzyme through gene expression and regulation by changing enzyme activity through posttranslational modifications, in particular phosphorylation. Gene expression is regulated by one or more transcription factors (TFs) and signaling molecules (Signals) regulate both transcription factor activity and metabolic enzyme activity by changing the phosphorylation status. This figure was modified from Supplementary Figure 1 of Kokaji et al. (2020).

**Fig. S2.**
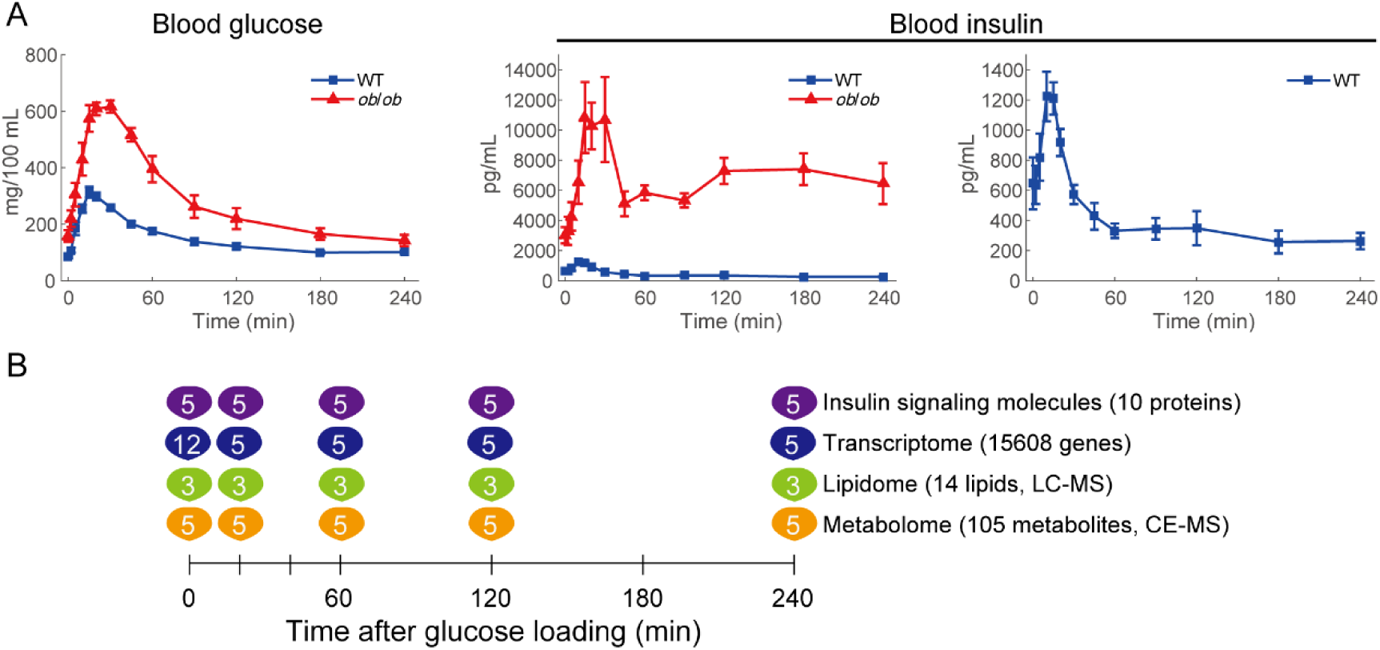
Oral glucose administration and multiomic measurements. (**A**) Blood glucose and blood insulin of WT mice (blue) and *ob*/*ob* mice (red) during oral glucose administration. The data of blood glucose and insulin levels measured in our previous study are shown (Kokaji et al., 2020).The means and SEMs of five mice per genotype are shown. (**B**) We orally administered glucose to 16 h-fasting WT and *ob*/*ob* mice, and collected the skeletal muscle at 0, 20, 60, 120, and 240 min after administration. We performed metabolomics, transcriptomics, and Western blotting for the phosphorylation of insulin signaling molecules in the skeletal muscle. The number of mice per genotype in each measurement is shown at each time point. This figure was modified from Supplementary Figure 2 of Kokaji et al. (2020).

**Fig. S3.**
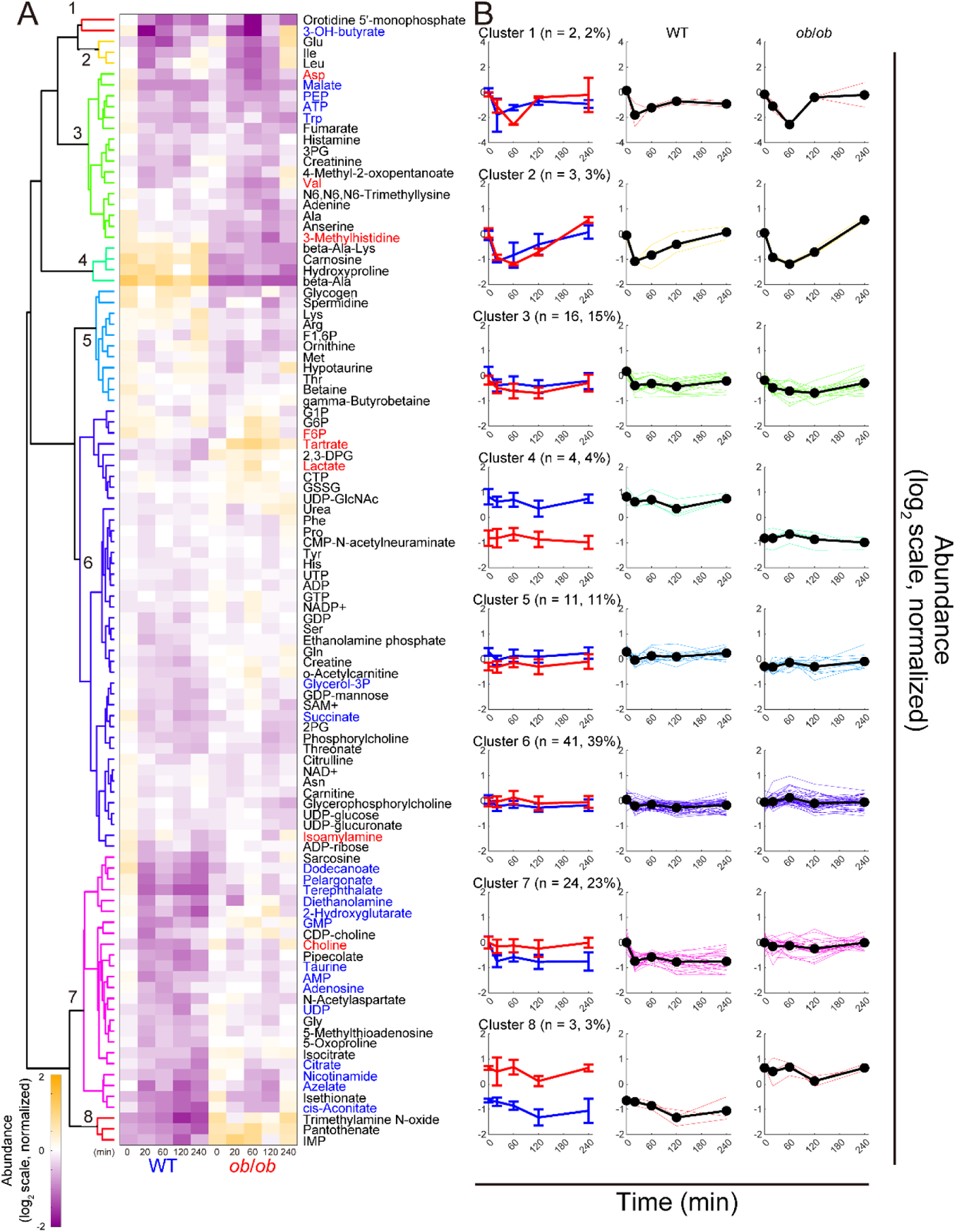
Hierarchical clustering of time courses of metabolites in the skeletal muscle. (**A**) The heat map and hierarchical clustering of the time courses of metabolites in the skeletal muscles of WT and *ob*/*ob* mice following oral glucose administration. The colors of and numbers on tree diagram indicate the cluster of each metabolite. To investigate the changes from fasting state, two time courses of each metabolite were divided by the geometric mean of the values of WT mice and *ob*/*ob* mice in fasting state (0 min), and then log_2_-transformed. The colors of the names of metabolites indicate WT mice-specific glucose-responsive metabolites (blue), *ob*/*ob* mice-specific glucose-responsive metabolites (red), and metabolites that are not glucose-responsive (black). (**B**) Averaged time courses of the metabolites for all eight clusters. Left panel shows averaged time courses of the metabolites as the mean and standard deviation in a cluster for WT mice (blue) and *ob*/*ob* mice (red). Middle panel (WT mice) and right panel (*ob*/*ob* mice) show average (thick line) and individual (thin line) time courses of the metabolites in a cluster in WT or *ob/ob* mice. Clusters 1, 2, and 3 was comprised of metabolites which were decreased in both WT and *ob/ob* mice, and the responses in cluster 1 were largest of the three clusters. Orotidine 5’-monophosphate and 3-OH-butyrate was classified in this cluster. The responses in cluster 2 were larger than those in cluster 3. Cluster 2 consisted of three amino acids; valine, leucine, and glutamate. Other amino acids (aspartate, valine, alanine, and tryptophan), downstream metabolites of the glycolytic pathway (3-phosphoglyceric acid [3PG] and phosphoenolpyruvate [PEP]) and metabolites of the TCA cycle (fumarate and malate) were classified into cluster 3. Cluster 4 included metabolites that were more abundant in WT mice at all timepoints. This cluster mainly comprised β-alanine, carnosine, a dipeptide of β-alanine and histidine, and β-alanine-lysine. Metabolites in cluster 5 were also more abundant in WT mice; however, the difference was smaller compared to cluster 4. This cluster mainly comprised amino acids such as lysine, arginine, threonine, and ornithine. Fructose 1,6-bisphosphate (F1,6BP) and glycogen was also classified into cluster 5. Metabolites in cluster 6 were observed at slightly higher levels in *ob/ob* mice, and showed almost no changes by glucose administration. Many amino acids, metabolites of the central carbon metabolism, and nucleic acids were classified into this cluster. Metabolites of the glycolytic pathway, such as glucose-1-phosphate (G1P), glucose-6-phosphate (G6P), fructose 6-phosphate (F6P), and lactate were also included. Among these, F6P and lactate were significantly increased only in *ob/ob* mice. Metabolites in cluster 7 tended to decrease specifically in WT mice (14/24 metabolites showed significant decreases). Metabolites of the TCA cycle, such as citrate, isocitrate, and cis-aconitate, and nucleic acids (adenosine monophosphate [AMP], guanosine monophosphate [GMP], and adenosine) were included in this cluster. Metabolites in cluster 8 (inosine monophosphate, pantothenate, and trimethylamine N-oxide) were abundant in *ob/ob* mice compared to WT mice, and the difference was quite large.

**Fig. S4.**
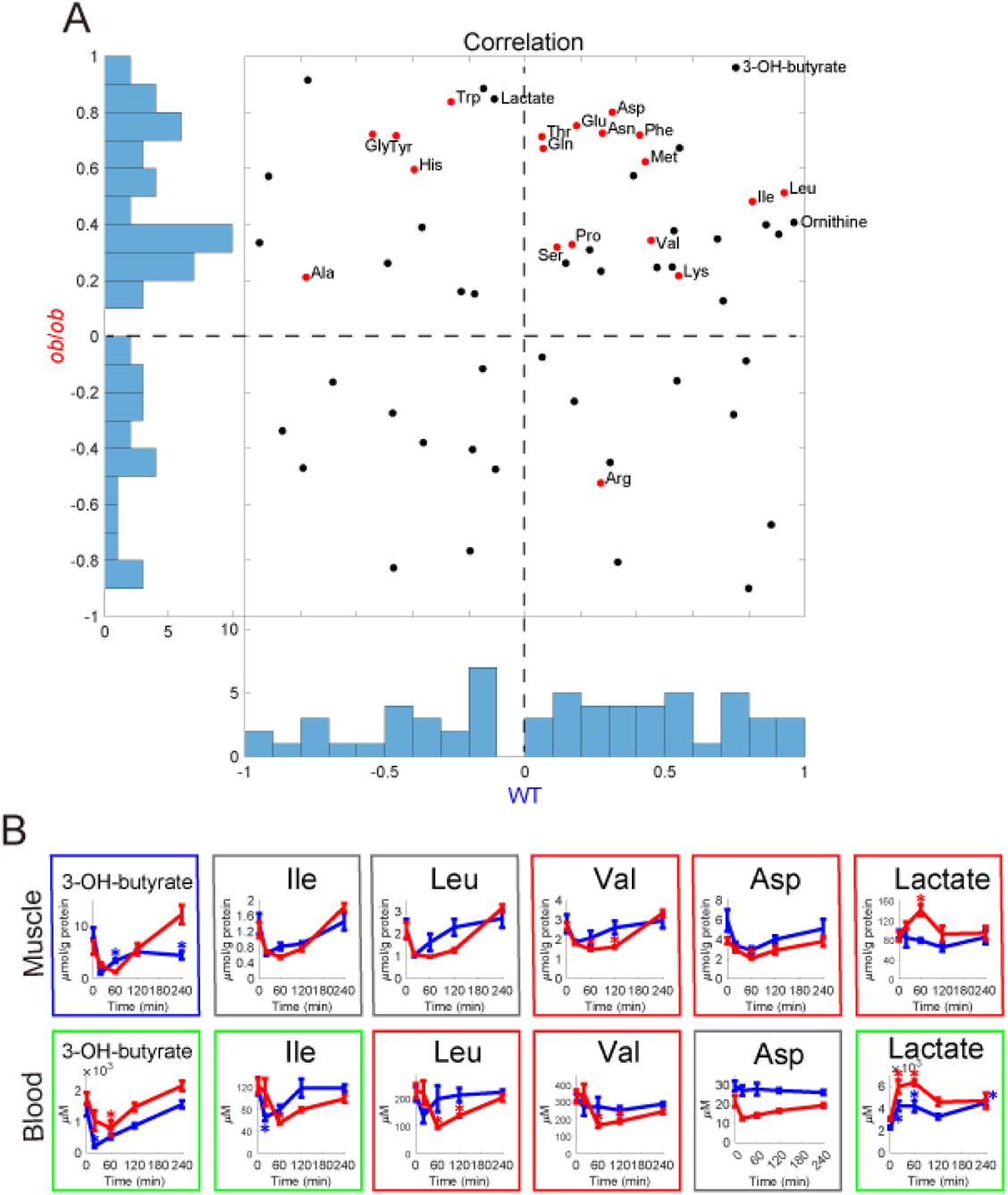
Time courses of metabolite changes in the skeletal muscle and in the blood. (**A**) Histograms and scatter plot of Pearson’s correlation coefficients between the time courses of changes in metabolites measured in skeletal muscle and blood in WT mice and *ob*/*ob* mice. Red dots indicate 19 proteogenic amino acids measured in both skeletal muscle and blood. (**B**) Time courses of changes in the indicated metabolites in the skeletal muscle and blood of WT mice (blue) and *ob*/*ob* mice (red) following oral glucose administration. The means and SEMs of five mice per genotype are shown. The colors of the frames indicate common glucose-responsive metabolites (green), WT-specific glucose-responsive metabolites (blue), *ob*/*ob*-specific glucose-responsive metabolites (red), and not glucose-responsive metabolites either in WT mice or in *ob*/*ob* mice (gray). *q value < 0.1 and absolute log_2_ fold change > 0.585.

**Fig. S5.**
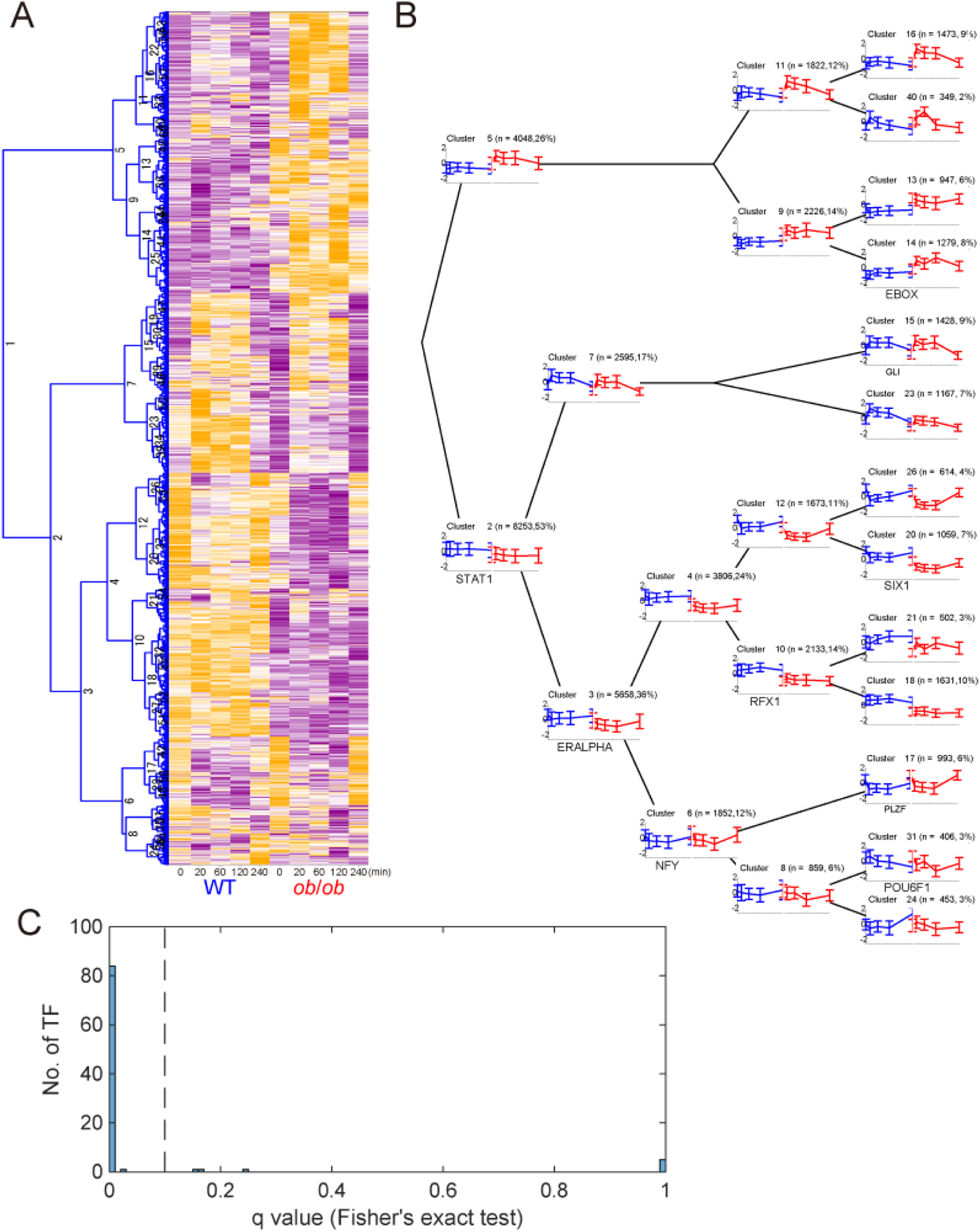
Hierarchical clustering of the time courses of gene expression in the skeletal muscle and inference of regulatory connections between transcription factors and genes. (**A**) The heat map and hierarchical clustering of the Z-score normalized time courses of gene expressions in the skeletal muscle of WT and *ob*/*ob* mice following oral glucose administration. The hierarchical clustering was performed using Euclidean distance and Ward’s method. The numbers on the tree diagram indicates the cluster identity. Each cluster includes only the genes that show a significant response at any time point either in WT mice or *ob*/*ob* mice or significant differences between WT mice and *ob*/*ob* mice before glucose administration (0 min). (**B**) The averaged time courses of the gene expression for each cluster of WT mice (blue) and *ob*/*ob* mice (red). The mean and standard deviation of the time courses of gene expressions in the cluster are shown. The time courses are presented on the tree diagram of hierarchical clustering. Significantly enriched transcription factor motifs (q value < 0.1) in the cluster are described with the time courses. According to the enriched transcription factor motifs, we defined the regulatory connections between the transcription factors and the genes in the cluster. To avoid overestimation, we excluded a cluster from the inference if the transcription factor binding motif was more enriched in the children clusters. The remaining transcription factor motifs, but not the excluded transcription factor motifs, are described here. The transcription factor motifs enriched in the upstream clusters are not described in the downstream clusters. (**C**) The histogram of the q values for the overlaps between the inferred genes of the transcription factors and those predicted from ChIP data. The ChIP data were obtained from the ChIP-Atlas database (Oki et al., 2018). The q values were calculated by the one-tailed Fisher’s exact test and Benjamini–Hochberg procedure (Yoav Benjamini, 1995).

**Fig. S6.**
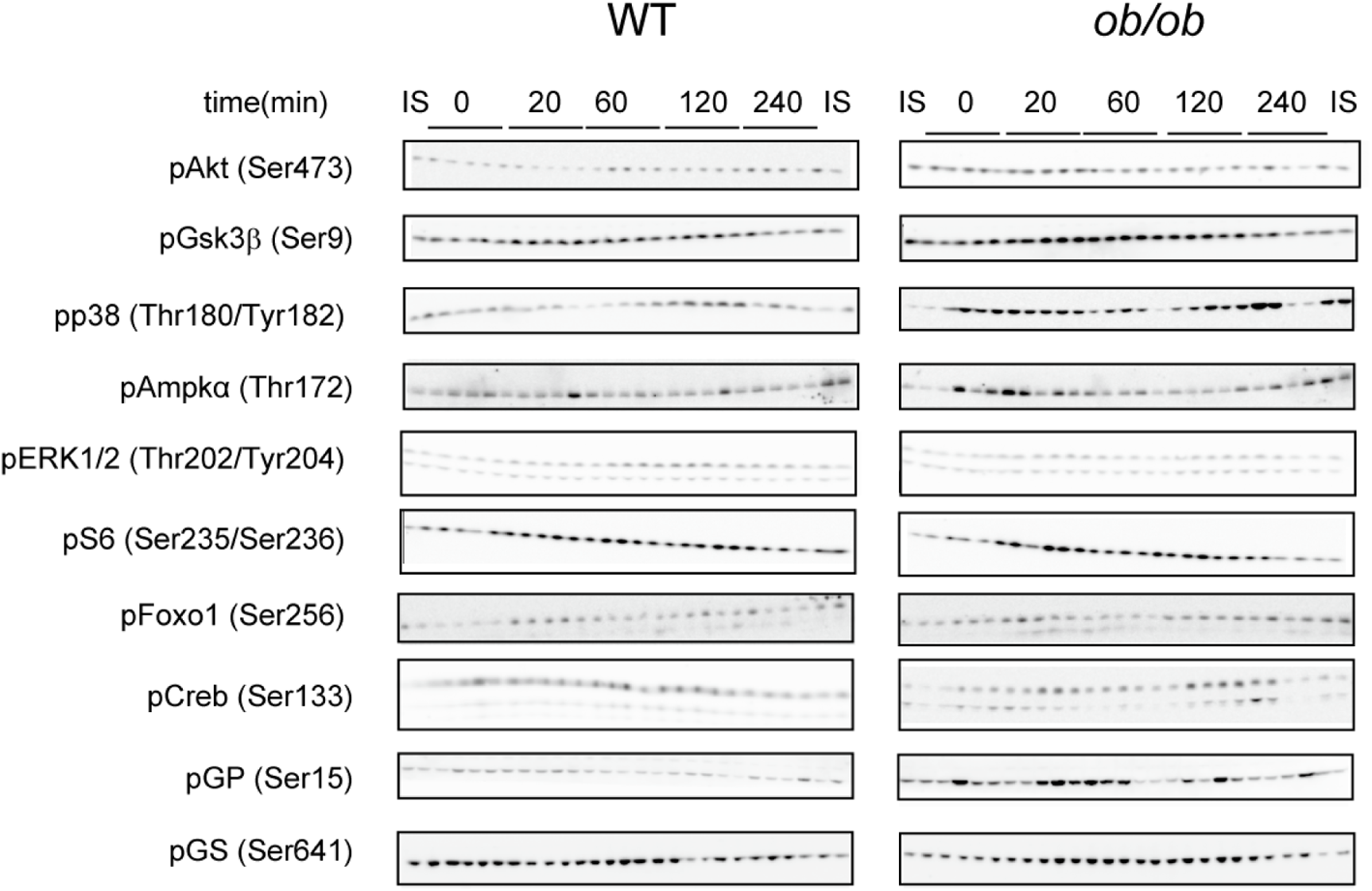
Western blotting for insulin signaling molecules. The phosphorylation of the indicated insulin signaling molecules in the skeletal muscle of WT and *ob*/*ob* mice at the indicated time point after oral glucose administration. Residues in parentheses indicate the phosphorylation site(s) (human sequence numbering) recognized by the antibodies. Western blot data for all mice are shown (n = 5 mice per genotype for glucose administration).

**Fig. S7.**
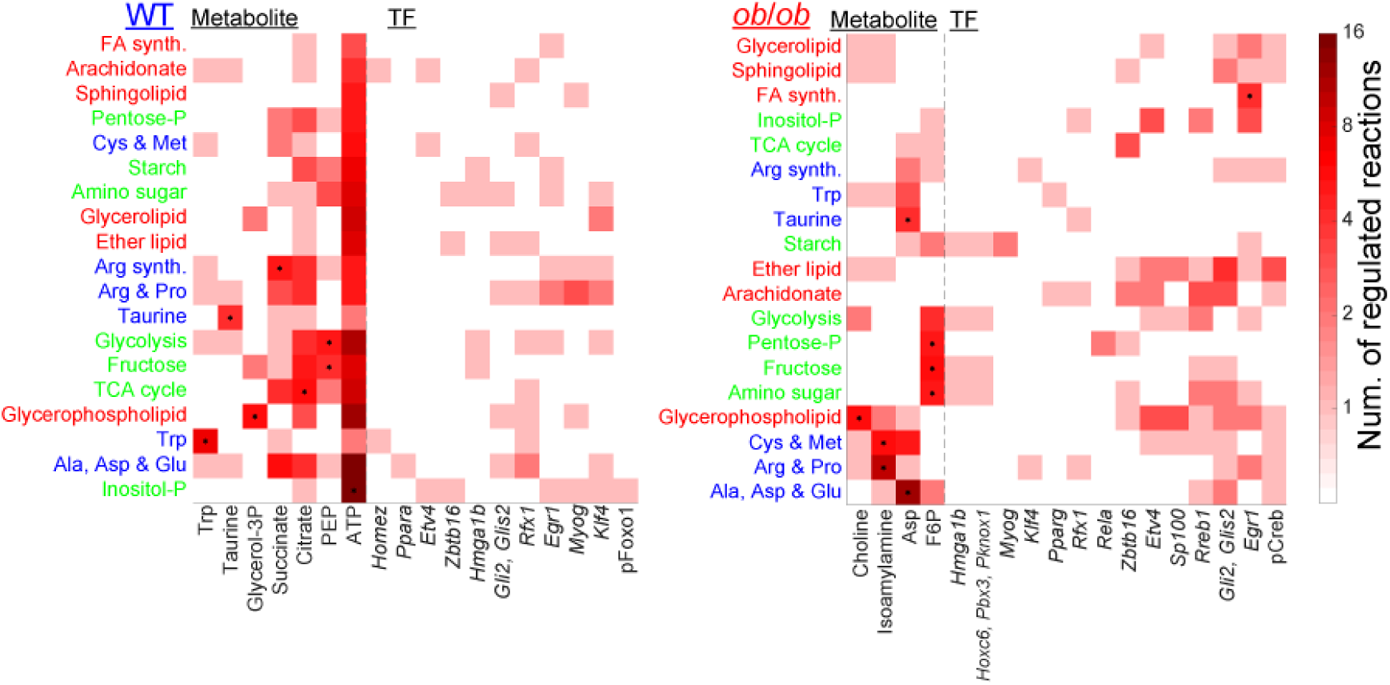
Metabolic reactions regulated by glucose-responsive molecules in each metabolic pathway node. Heat maps showing the number of regulated metabolic reactions in each metabolic pathway node (rows) by each glucose-responsive metabolite (left columns) and each transcription factor-dependent glucose-responsive genes of metabolic enzymes (right columns) in WT and *ob/ob* mice. The * symbols indicate significant associations (q value < 0.01) between metabolic reactions in the metabolic pathway node and those regulated by glucose-responsive molecules (Data File S10). The q values were calculated by the one-tailed Fisher’s exact test and Benjamini–Hochberg procedure (Yoav Benjamini, 1995). Only metabolic pathway nodes with significant associations with any glucose-responsive molecule are shown. Only glucose-responsive metabolites with significant associations with any metabolic pathway node are shown.

Data File S1. Metabolomic data.

Data File S2. Lipidomic data.

Data File S3. Transcriptomic data.

Data File S4. Pathway enrichment analysis of glucose-responsive genes.

Data File S5. Enrichment analysis of gene clusters.

Data File S6. Inferred regulatory connections between transcription factors and genes.

Data File S8. Western blotting data.

Data File S9. Regulatory transomic network for glucose-responsive metabolic reactions.

## Acknowledgements

We thank Maki Ohishi, Ayano Ueno, Hiroko Maki, Keiko Endo, and Sanae Ashitani (Keio University) for their technical assistance with metabolomic analysis using CE–MS; and our laboratory members for critically reading this manuscript and for their technical assistance with the experiments. The computational analysis of this work was performed in part with support of the super computer system of the National Institute of Genetics (NIG), Research Organization of Information and Systems (ROIS).

## Author contributions

T.K., M.E., A. Hatano, K.M., Y.I., R.E., and H.K. designed and performed the animal experiments, enzyme assays and western blot analysis. A. Hirayama and T.S. performed metabolomic analysis using CE–MS. Y.S. performed RNA sequencing transcriptomic analysis. T.K., K.Y., S.O., M.F., K.H., Y.I., S.U., A.T., Y.P., H.M., D.L., Y.B., T.T., and H.O. performed transomic analysis. The writing group consisted of T.K., M.E., H.I., and S.K. The study was conceived and supervised by T.K. and S.K.

## Data and materials availability

Sequencing data measured in this study have been deposited in the DNA Data Bank of Japan Sequence Read Archive (DRA) (www.ddbj.nig.ac.jp/) under the accession no. DRA010972 and DRA013659. All other data needed to evaluate the conclusions in the paper are present in the paper or Supplementary Materials. The code used for the analysis in this paper is available upon request.

